# Antimicrobial cetylpyridinium chloride suppresses mast cell function by targeting tyrosine phosphorylation of Syk kinase

**DOI:** 10.1101/2024.07.04.602096

**Authors:** Bright Obeng, Lucas J. Bennett, Bailey E. West, Dylan J. Wagner, Patrick J. Fleming, Morgan N. Tasker, Madeleine K. Lorenger, Dorothy R. Smith, Tetiana Systuk, Sydni M. Plummer, Jeongwon Eom, Marissa D. Paine, Collin T. Frangos, Michael P. Wilczek, Juyoung K. Shim, Melissa S. Maginnis, Julie A. Gosse

## Abstract

Cetylpyridinium chloride (CPC) is a quaternary ammonium antimicrobial used in numerous personal care products, human food, cosmetic products, and cleaning solutions. Yet, there is minimal published data on CPC effects on eukaryotes, immune signaling, and human health. Previously, we showed that low-micromolar CPC inhibits rat mast cell function by inhibiting antigen (Ag)-stimulated Ca^2+^ mobilization, microtubule polymerization, and degranulation. In this study, we extend the findings to human mast cells (LAD2) and present data indicating that CPC’s mechanism of action centers on its positively-charged quaternary nitrogen in its pyridinium headgroup. CPC’s inhibitory effect is independent of signaling platform receptor architecture. Tyrosine phosphorylation events are a trigger of Ca^2+^ mobilization necessary for degranulation. CPC inhibits global tyrosine phosphorylation in Ag-stimulated mast cells. Specifically, CPC inhibits tyrosine phosphorylation of specific key players Syk kinase and LAT, a substrate of Syk. In contrast, CPC does not affect Lyn kinase phosphorylation. Thus, CPC’s root mechanism is electrostatic disruption of particular tyrosine phosphorylation events essential for signaling. This work outlines the biochemical mechanisms underlying the effects of CPC on immune signaling and allows the prediction of CPC effects on cell types, like T cells, that share similar signaling elements.

## Introduction

Cetylpyridinium chloride (CPC) is a widely used cationic quaternary ammonium^1^ (structure in **Fig. 4A**) antimicrobial agent. While the FDA effectively banned CPC from hand soaps and sanitizers^2,3^, it remains at high concentrations (up to thousands of micromolar) in many consumer products such as mouthwashes, lozenges, lip balms, household cleaning solutions, deodorants, and hair styling products^1,4,5^. It is used to treat gingivitis^6^ and halitosis^7^ and in general as an antibacterial. CPC is also employed in the agricultural sector for food processing, particularly poultry^8,9^. CPC has also been investigated in lab and clinical studies for efficacy against certain viruses (as previously summarized^10^, including SARS-CoV-2^11^). The widespread usage of CPC underscores the importance of understanding its toxicology, as people are exposed to high concentrations of CPC via diverse routes of application. However, very little information is published regarding CPC’s effect on immune signaling and function.

CPC is retained in human tissues^12^. A recent study in mice shows that CPC crosses the blood-brain barrier and is toxic to oligodendrocytes^13^. Based upon previous literature reports regarding the bioavailability of CPC^14–16^, exposure estimates from the European Union Scientific Committee on Consumer Safety^4^, along with our literature review and calculations^10^, it is possible that human blood levels of CPC may average ∼0.3 μM from personal care product usage with an additional ∼0.3 μM from consumption of CPC-treated chicken. However, to our knowledge, no direct measurements (beyond the 1978 saliva study^12^) of the human body burden of CPC have been published.

Experiments in the current study were performed at low-micromolar CPC doses. The CPC doses used in this study are not cytotoxic, as determined by lactate dehydrogenase and trypan blue assays^17^, a plasma membrane (PM) potential determination^18^, and another fluorescence-based cytotoxicity test^10^. Also, while CPC acts as a cell-lysing detergent when present at high concentrations, above its critical micelle concentration (CMC) of 600 - 900 μM^19–21^, all doses in the current study are ∼100-fold lower than the range in which CPC can detergent-lyse cells.

Hexadecylbenzene (HDB) is chemically structurally similar to CPC^22^, apart from its positively-charged nitrogen in the head group ring, as shown in **Fig. 4A**. Thus, the relevance of CPC’s quaternary nitrogen and positive charge can be isolated and compared with the other structural features (saturated lipid tail and conjugated ring) by comparing CPC and HDB effects.

Mast cells (MCs) are found in most tissues^23,24^ and play key roles in the immune^25,26^ and nervous system functions^27,28^ via cytokine release and a physiological process called degranulation, in which they release highly bioactive mediators such as histamine and serotonin. Previously, we showed that CPC inhibits degranulation of rat mast cells in a time- and dose-dependent manner^17,18^. MCs are enriched at environmental interfaces (such as skin, oral/ respiratory/ gastrointestinal mucosa^27^, nerve terminals^23,24^); thus, they are poised for exposure to CPC-containing products via various routes.

In this study, the LAD2 human and the RBL-2H3 (rat basophilic leukemia cells, clone 2H3; “RBL”) MC models are employed. The LAD2 cells express ultrastructural features of mature human MC^29^. RBL cells are homologous to mature human MCs, rodent mucosal MCs, and basophils regarding functionality and signaling machinery^30–33^. RBL cells respond to external/environmental stimuli similarly to primary bone marrow-derived mast cells^34–36^. Thus, LAD2 and RBLs are scientifically suitable for toxicological/pharmacological mast cell signal transduction studies.

The high-affinity IgE receptor FcεRI^27,37^ on the surface of MCs binds IgE^38^, which is aggregated by multivalent antigen (Ag, or allergen) or other crosslinkers, leading to a signaling cascade. Streptavidin, dinitrophenyl-bovine serum albumin (DNP-BSA, a multivalent Ag), and bivalent anti-IgE IgG (see schematic in **Fig. 3**) are examples of IgE-FcεRI crosslinkers. A multivalent (Ag) can crosslink more than two IgE-FcεRI receptor complexes at a time whereas the bivalent anti-IgE IgG can crosslink only two IgE-bound FcεRI receptors (see schematic in **Fig. 3**).

Upon Ag crosslinking of IgE-FcεRI, conserved immunoreceptor tyrosine-based activation motifs (ITAMs) in the cytoplasmic tails of the high-affinity IgE receptor FcεRI β and γ subunits^39,40^ become phosphorylated by the Src family Tyr kinase Lyn^40,41^, which is anchored to the PM^42,43^. Once phosphorylated, the anionic phosphorylated tyrosines within the ITAMs function as a docking and activation site for proteins with cationic Src-homology-2 (SH2) domains^42,44,45^. All SH2 domains contain a positively-charged binding site that utilizes Arg to bind to negatively-charged phosphorylated Tyr^46^.

Lyn kinase exists as two isoforms, A and B^47^. The sequence of Lyn is highly conserved between the isoforms apart from 21 amino acids near the N terminus missing from isoform B^48^. Connected to the PM via a saturated 14-carbon myristoyl and 16-carbon palmitoyl anchors, Lyn also contains an SH2 domain^49^, along with positive^50–52^- and negative-regulating^53^ phosphotyrosine residues, Tyr397 (Y396 or Y397) and Tyr507 (Y507 or Y508), respectively (**Fig. 1**). In literature, Lyn Y507/508 are synonymous^53^, similar to Lyn Y396/397. Prior to Ag stimulation, Lyn is inactive but remains constitutively phosphorylated^41,54^ (see **Fig. 9C**), including at Y507 which negatively regulates mast cell degranulation. Lyn activation begins with dephosphorylation of Y507^55,56^. Lyn Y397, found in the activation loop, regulates degranulation positively. Autophosphorylation and activation occur at the Y397 site, causing Lyn to possess its greatest enzymatic activity^50–52^. Lyn exists in a closed conformation when inactive and transitions to an open conformation upon activation^57^ (**Fig. 1**).

**Fig. 1.**
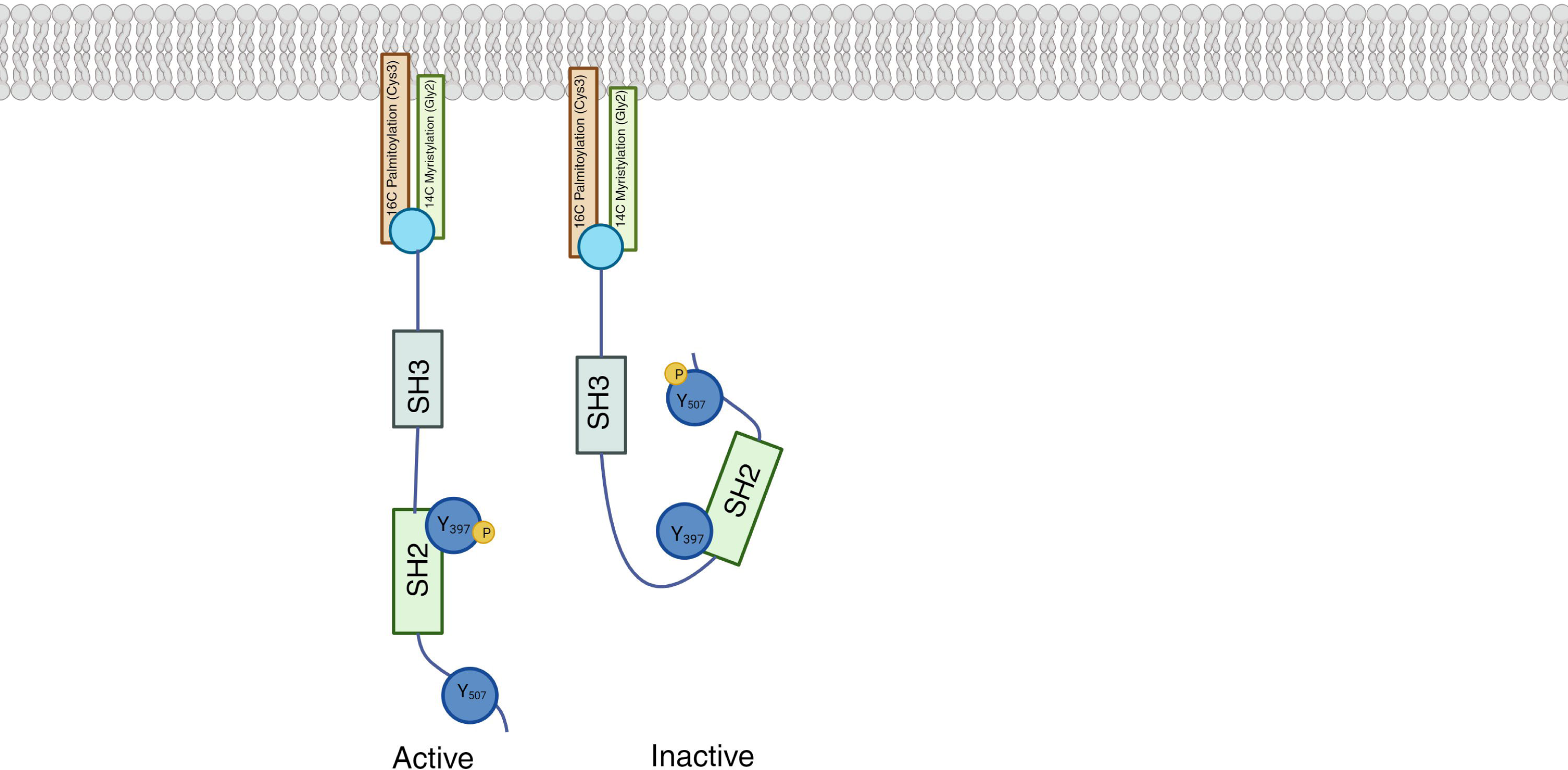
Schematic representation of regulating tyrosine residues in Lyn kinase. Lyn is equipped with a myristoylation and palmitoylation site, an SH2 and SH3 domain, a positively regulating Tyr^397^, and a negatively regulating Tyr^507^ (Y507). Lyn activity is determined by the conformation of the kinase. The phosphorylation state of tyrosine residues Tyr^397^ and Tyr^507^ indicates conformation.

Phosphorylated FcεRIγ recruits spleen tyrosine kinase (Syk), which contains SH2^44,45^ and is phosphorylated by Lyn^58–60^. Syk plays an essential role in MCs^61,62^, activating adaptor molecules like LAT^63^, which is necessary for degranulation^63–67^. Syk-phosphorylated LAT is a scaffold for phospholipase C gamma (PLCγ)1 and PLCγ2^63^. LAT-recruited PLCγ1 is then activated by Syk^68^.

PLCγ1 binds phosphatidylinositol 4,5-bisphosphate (PIP_2_) in the PM inner leaflet via its Pleckstrin Homology (PH) domain^69^ and catalyzes PIP_2_ hydrolysis to generate inositol-1,4,5-trisphosphate (IP_3_) and diacylglycerol (DAG). IP_3_ initiates Ca^2+^ mobilization by binding to its receptors^70^ on the endoplasmic reticulum (ER) membrane, thus inducing ER Ca^2+^ efflux^71^, thus inducing store-operated Ca^2+^ entry into the PM^72–74^, required for degranulation^75^. Together, elevated cytosolic Ca^2+^ and DAG^76–78^ activate enzymes needed for degranulation^79–82^. In activated MC, microtubules polymerize in a Ca^2+^-dependent manner^83–87^, then serve as the “railroad tracks” along which granules are transported to the PM for degranulation^83^ . At the PM, granules interact with SNAREs for exocytosis^88–92^, releasing bioactive mediators^93^, which include serotonin, histamine, and beta-hexosaminidase^94^.

Previously, we reported CPC suppression of Ag-stimulated Ca^2+^ efflux from the ER and of subsequent cytosolic Ca^2+^ rise (via PM CRAC channel activation) necessary for the polymerization of microtubules^18^. Thus, a target upstream of IP_3_ generation/ER Ca^2+^ mobilization is responsible for the CPC inhibition of degranulation. Therefore, the current study examined whether CPC affects early tyrosine phosphorylation events that precede and trigger Ca^2+^ mobilization in MCs. We hypothesized that the positively-charged CPC electrostatically interferes with the tyrosine phosphorylation of kinases, explaining the inhibition of Ca^2+^ mobilization and degranulation. We investigated the effects of CPC on global and specific tyrosine phosphorylation events in Ag-stimulated mast cells. We show that CPC inhibits early tyrosine phosphorylation events, particularly Syk kinase.

## Materials and Methods

### Cetylpyridinium chloride monohydrate preparation

Cetylpyridinium chloride (CPC; 99.7% purity, MP Biomedicals; CAS no. 6004-24-6) was prepared fresh on each experimental day as described^17^ in aqueous buffer without organic solvent. UV-Visible spectrophotometry (Shimadzu UV-1900i) was used to confirm CPC concentration^4,17^; nominal concentrations calculated from grams added and buffer volume were close to the measured UV-Vis concentrations. Bovine serum albumin (BSA; 98% purity, Millipore Sigma; CAS no. 9048-46-8) was added to diluted CPC solutions at 1 mg/mL, resulting in BSA-Tyrodes (BT)^95^ CPC-containing solutions. See the schematic on each figure for the CPC treatment conditions. Doses of CPC and treatment conditions used in this study are not lethal to cells^17,18^, as measured by several endpoints. In all experiments in which CPC was used, BT was the vehicle^17,18^.

### Hexadecylbenzene preparation

Hexadecylbenzene (HDB; 99+% purity, stock stored in a desiccator at 4 °C, TCI America™; CAS no. 1459-09-2) was prepared on each experimental day in Tyrodes buffer^95^ after first dissolving HDB in 100% dimethyl sulfoxide (DMSO; 99.7+% purity, Sigma Aldrich; CAS no. 67-68-5). Note that the melting temperature of HDB is close to room temperature (RT)^96^, so crystals may appear to melt or become “watery” while working with pure HDB. Care was taken throughout procedures to protect this light-sensitive chemical. Firstly, HDB, 0.14 g, was dissolved in 2 mL of prewarmed, 37 °C, 100% DMSO (to a nominal concentration of 0.231 M) by vortexing, followed by sonication (Branson 1200 ultrasonic cleaner; Branson Ultrasonics, Danbury, CT, USA) at 40 °C for 10 min. Following sonication and through the rest of the procedure, the solution appeared well-dissolved, without particulate matter or precipitate. Next, 100 μL of Tyrode buffer was removed from 50 mL pre-warmed Tyrodes buffer, and 100 μL of the dissolved HDB-DMSO was added (to a nominal concentration of 462 µM) with vortexing, then sonicated at 40 °C for 20 min. After sonication, 30 ml of sonicated HDB-DMSO solution was further diluted via addition to 100 ml of prewarmed Tyrodes buffer (to a nominal concentration of 106.6 µM), followed by several min of stirring on a magnetic stir plate. HDB concentrations were determined using UV-Vis spectrophotometry (Shimadzu, UV-1900i) (see **Fig. S1**) and the Beer-Lambert equation (A_262_ = εlc); published data in an undefined solvent show HDB absorption maximal at 262 nm^96^. The UV-Vis spectrum was measured from 255 nm (not lower due to absorption of the cuvettes utilized), to 320 nm. A control, DMSO in Tyrodes buffer, which was prepared in tandem through all steps including sonications and various dilutions, was used as the UV-Vis blank. Because an extinction coefficient for HDB dissolved in aqueous buffer (with DMSO vehicle) was not located in the literature, we used the ε_260_ of CPC, 4389 M^−1^ cm^-1^ ^4^ to estimate HDB concentration. Bovine serum albumin (BSA; 98% purity, Millipore Sigma; CAS no. 9048-46-8) was added at 1 mg/mL to create a final HDB solution in BT. The final DMSO concentration used in the experiments was 0.046% v/v; this concentration does not affect ATP levels, degranulation, or cytotoxic of RBL mast cells^97^.

## Cell culture

### RBL-2H3 mast cells

RBL mast cells were cultured as described ^95^. Mycoplasma tests were performed to ensure cells were free of contamination (data not shown).

Cells were sensitized with anti-dinitrophenyl (DNP) mouse IgE as published^18^. IgE-bound receptors were crosslinked in most experiments with multivalent dinitrophenyl-bovine serum albumin (DNP-BSA)^18^, which crosslinks more than two IgE-bound FcεRI receptors at a time (see schematic in **Fig. 3**). An alternate stimulator, the bivalent crosslinker anti-Mouse IgG (Fab specific) antibody produced in goat (Millipore Sigma), that crosslinks only two IgE-bound FcεRI receptors at a time (see schematic in **Fig. 3**), was used to generate the data in **Fig. 3**.

### LAD2 cells

LAD2 cells, which are human mast cells that grow in suspension, were cultured as described^98^. Mycoplasma tests were performed to ensure cells were free of contamination (data not shown). To stimulate the cells, cells were sensitized with biotinylated human IgE (0.1 µg/mL) (Enzo Life Sciences) for 24 h^99^ at 37 °CO_2_, and excess IgE was washed off with BT before stimulation with streptavidin (Millipore Sigma; CAS no. 9013-20-1).

### Degranulation assay

RBL-2H3 mast cell degranulation responses were measured as described ^18^. Schematics in Figures 3 and 4 depict CPC and HDB treatment conditions for the degranulation responses, respectively.

LAD2 mast cell degranulation responses were measured using a fluorescence-based assay as previously described^100^, adapted with modifications for use in LAD2 cells^98^. LAD2 mast cells, 2*10^6^ in 4 mL total, were sensitized with biotinylated human IgE (0.1 µg/mL) for 24 h at 37 °C/5% CO_2_ in a 24-well sterile plate. Following sensitization, cells were centrifuged at 450 x g at RT for 5 min, supernatant was removed and discarded, washed with BT (1 mL per 1*10^6^ cells) to remove unbound IgE, and again centrifuged at 450 x g at RT for 5 min. After supernatant removal, cells are reconstituted gently in BT and then plated at a density of 100,000 cells/well (in 50 µL of BT) and pretreated with CPC or BT for 1 h or for 30 min (see schematic in **Fig. 2**) by adding 50 µL of 2X CPC solutions and incubating at 37 °C/5% CO_2_. After the pretreatment, without discarding the solutions, cells are stimulated with 100 µL of 2X biotinylated-streptavidin ± 1X CPC for 30 mins at 37 C/5% CO_2_. Spontaneous (no stimulation), Background (no cells), and Triton-X 100 (maximal granule release) samples were included as noted^100^. Following stimulation, cells in the 96-well plate are spun at 450 x g for 5 min^98^. After spinning down the cells, the plate is immediately put on ice, and the beta-hexosaminidase contained in the supernatant is measured as described^100^.

**Fig 2.**
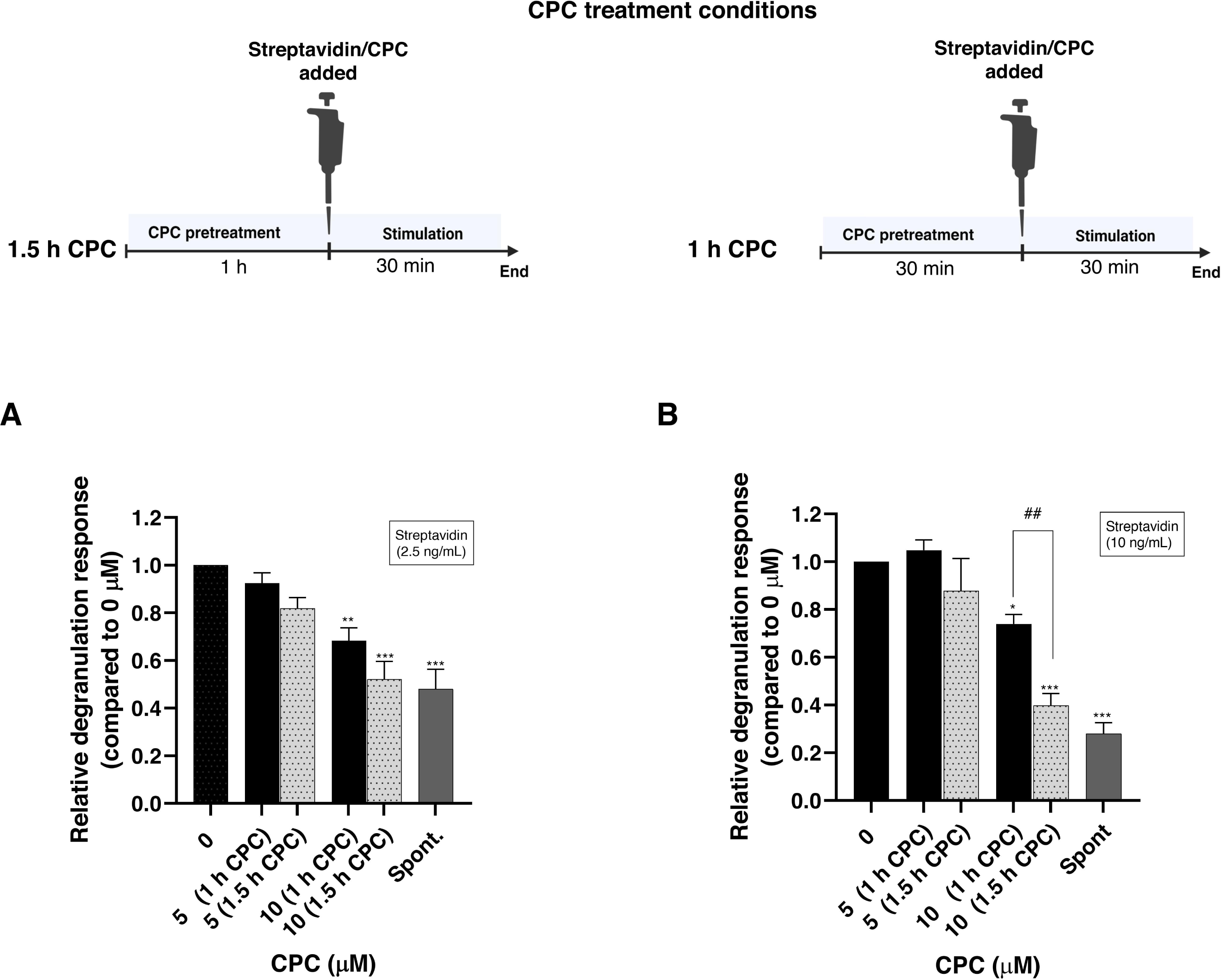
Relative degranulation response of LAD2 human mast cells exposed to non-cytotoxic doses of CPC. Cells were sensitized with biotinylated human IgE for 24 hours. For spontaneous release (“Spont.” on x-axes), cells were exposed to BT for 30 min without streptavidin. Samples denoted “1.5 h CPC” (grey bars; see treatment schematic) had CPC pretreatment for 1 h before the 30 min streptavidin/CPC co-exposure. Samples denoted “1 h CPC” (black bars; see treatment schematic) underwent 30 minutes CPC pretreatment before the 30 min streptavidin/CPC co-exposure. Cells were stimulated with 2.5 **(A)** or 10 ng/mL **(B)** streptavidin for 30 min. Values presented are means normalized to the control, ± SEM, of at least 3 independent days of experiments; each cell treatment was performed in duplicate. Statistically significant results in comparison to the 0 μM control of each graph are represented by *p *<* 0.05, **p *<* 0.01, and ***p *<* 0.001, ****p < 0.0001 or in comparison to particular samples are indicated by vertical lines are represented by ##p *<* 0.01, each determined by one-way ANOVA followed by Tukey’s post-hoc test.

### In-Cell Western Assay

Published In-Cell Western (ICW) procedures^101^ were adapted for use with RBL-2H3 cells with CPC treatment. RBL-2H3 cells were plated in a 96-well plate at 75,000 cells/well in RBL media (150 µl/well) and incubated overnight at 37 °C/5% CO_2_. Media was removed with an aspirator, and cells were sensitized with anti-DNP mouse IgE (0.1 µg/mL; 100 µL/well) (Millipore Sigma) for 1 hour at 37 °C/5% CO_2_. Following sensitization, media was removed, cells were washed with BT twice (200 µL/well/wash), washes were aspirated, and cells were pre-treated with BT or 10 µM CPC (200 µL/well) for 30 min at 37 °C/5% CO_2_. At the end of the pre-treatment, liquid was removed from wells with an aspirator, the plate was placed on a plate heater, and cell groups “Spont.,” “Ag,” and “Ag + CPC” were exposed to 200 µL/well of BT, Ag (0.0007 µg/mL), or Ag (0.0007 µg/mL) + CPC (10 µM), respectively, for 5 min at 37 °C/5% CO_2_. Cells were then aspirated, washed once with PBS (200 µL/well) with Combitip, aspirated, and fixed with 150 µL/well Fisher Science Education™ Formaldehyde Solution (Thermo Fisher Scientific; CAS no. 50-00-0, 67-56-1, 7732-18-5) diluted to 3.7% in phosphate buffered saline (PBS; Millipore Sigma) for 20 min at RT. Next, fixed cells were aspirated, washed twice with PBS (200 µL/well) with Combitip, aspirated, and permeabilized with 200 µL/well of RT 0.1% Triton X-100 (Thermo Fisher Scientific, CAS no. 9002-93-1) in PBS for 5 min on a plate shaker. The permeabilization process was repeated three additional times. Cells were then blocked with 150 µL/well Intercept(R) PBS Blocking Buffer (LI-COR) overnight at 4 °C with no shaking^102^. Cells were then incubated with the anti-phosphotyrosine primary antibody (Ab) P-Tyr-1000 MultiMab™ Rabbit mAb mix (Cell Signaling Technology) at 1:200 in Intercept(R) PBS Blocking Buffer (LI-COR) for 2 h at RT with gentle shaking. The following day, cells were washed twice with PBS + 0.1% Tween 20 (Millipore Sigma; CAS no. 9005-64-5) for 5 min with gentle shaking. Cells were then incubated with gentle shaking (and protected from light) for 1 h with 50 µL/well of the following mixture: IRDye^®^ 800CW Goat anti-Rabbit IgG Secondary Ab (LI-COR) at 1:5000 plus CellTag™ 700 stain (for normalization; LI-COR) at 1:1000, diluted in Intercept(R) Blocking Buffer with 0.2% Tween 20. Concurrently with this whole process, “secondary antibody-alone” samples were included–these cells received no IgE/Ag/CPC and no primary Ab but only secondary Ab, in order to control for non-specific binding of the secondary Ab to cells. Following washing two times for 5 min at RT with PBS + 0.1% Tween 20, the plate was imaged using the LI-COR Odyssey CLx Infrared Imaging system at 700 and 800 nm with a 42 µm resolution, medium intensity, 3.0 mm focus offset. Following scanning, the 700 and 800 nm channels were aligned using the Image Studio software (version 5.2). The signal in the 800 nm channel represents the tyrosine phosphorylation level, and the signal in the 700 channel is used for normalization. Values from the 800 nm channel of “secondary antibody-alone” wells were subtracted from the experimental values in the 800 channel. Next, the ratio of these background-subtracted secondary Ab (800) sample values to each corresponding CellTag (700) value (from the same well) was calculated. Finally, these ratios were normalized to spontaneous samples (no Ag, no CPC).

To assess the CPC effect on constitutive tyrosine phosphorylation by ICW (**Fig. S2**), cells were pre-treated with CPC (Spont. + CPC) or BT (for Spont.) for 30 min and given no Ag. There was no effect of CPC on constitutive phosphorylation.

### Western Blot (WB)

RBL-2H3 cells were plated at 0.5*10^6^ cells/well in 1.2 mL of RBL media on a 24-well, clear, TC-treated plate (Corning Costar(R)) and incubated overnight at 37°C/5% CO_2_. Cells were sensitized with anti-DNP mouse IgE (0.1 µg/mL) (Millipore Sigma), 400 µL/well, for 1 hour at 37 °C/5% CO_2_. Following sensitization, media was removed, cells were washed twice with warm BT (500 µL/well/wash), and then cells were pre-treated with BT or 10 µM CPC (for Spont. and Ag), 1 mL/well, for 30 min at 37°C/5% CO_2_. At the end of the pre-treatment, media were discarded. Immediately, cell groups “Spont.,” “Ag,” and “Ag + CPC” were stimulated with 1 mL/well of BT, Ag (0.0007 µg/mL), or Ag (0.0007 µg/mL) + CPC (10 µM), respectively, for 5 min or 2 min. This stimulation was performed with the bottoms of the wells in a shallow, pre-warmed 37 °C water bath in an incubator. Following stimulation, media were rapidly discarded, cells were placed on ice and immediately harvested with cold 2x Laemmli sample buffer (Bio-Rad), 200 µL/well, containing 5% β-mercaptoethanol (MP Biomedicals; CAS no. 60-24-2). Cells were harvested by pipetting, and lysates were placed into 1.5 mL screw-cap microcentrifuge tubes. These whole-cell lysates were then boiled for 5 min at 100 °C and immediately centrifuged at 4°C and 17,900 x g (Eppendorf, Centrifuge 5417R) for 10 min. Supernatants (100 µL) from the very top of each sample were removed to pre-chilled 1.5 mL tubes, stored on ice.

Protein concentrations for each sample were determined using the RC DC^TM^ Protein Assay (Bio-Rad) in sterile, flat-bottom 96 well plates (Greiner). The RC DC protein assay is compatible with reducing agents, detergents, and Laemmli buffer. The RC DC^TM^ protein assay was performed following the manufacturers’ instructions, except that the BSA (Bio-Rad) standard was dissolved in 5% H_2_O / 95% 2x Laemmli buffer (Bio-Rad) in order to simulate the cell lysate samples to be analyzed. Results were measured as absorbance at 750 nm. CPC treatment before and during Ag stimulation does not affect the total amount of protein in the whole cell lysate (**Fig. S3**).

An equal amount of protein, 12 μg (in a maximal volume of 15 μL) from each sample was loaded alongside a ladder (Chameleon^®^ Duo Pre-stained Protein Ladder, LI-COR) on 12% Mini-PROTEAN TGX Precast Protein gel, and proteins were separated by SDS-PAGE in TRIS-glycine-SDS running buffer, pH 8.3 running buffer (Thermo Fisher Scientific) with a Mini-Protean Tetra System (Bio-Rad). Proteins were transferred to a nitrocellulose membrane (pore size 0.45 µm; Bio-Rad) at 4°C overnight (16 hours, 30 V) in NuPAGE Transfer Buffer (Thermo Fisher Scientific) including 20% methanol (Thermo Fisher Scientific, CAS no. 67-66-1). Following the transfer, the membrane was dried for an hour (on Extra Thick Blot filter paper, Bio-Rad) on the benchtop at RT) to enhance protein retention on the membrane. Next, the blot was re-hydrated in TBS (Tris Buffered Saline, Millipore Sigma) for 5 min at RT with gentle agitation. After rinsing in MilliQ water, total protein was stained using Revert^TM^ 700 Total Protein Stain (LI-COR), imaged, and then destained according to manufacturer’s instructions. Total Protein Stain is recommended as the preferred normalization method for quantitative Western blotting ^103,104^. For all proteins except for total Syk, the membrane was blocked with 5% nonfat dry milk (Cell Signaling Technology) in TBS for 3 h, protected from light, at RT. After blocking, the membrane was washed three times, 5-10 min each, with 0.05% TBST (TBS + Tween20 [Millipore Sigma; CAS no. 9005-64-5]).

To detect tyrosine phosphorylation of proteins from whole cell lysates, the blot was probed with anti-phosphotyrosine monoclonal Ab clone 4G10 from mouse (Millipore Sigma) (1:5000, diluted in 1% nonfat dry milk-TBS) by incubating overnight at 4 °C with gentle shaking and protection from light. Next, three washes with TBST (10 min/wash) were performed. The anti-phosphotyrosine Ab 4G10 was detected with IRDye 800CW goat anti-mouse (LI-COR) secondary antibody (1:20000, diluted in 1% nonfat dry milk-TBS) by incubating at RT for 3 h, protected from light.

To detect phosphorylation of the Y397 residue of Lyn, the blot was probed with anti-phospho-LYN (Tyr397) monoclonal Ab from rabbit (Cell Signaling Technology, [1:1000 diluted in 1% nonfat dry milk-TBS]) by incubating at 4°C overnight with gentle shaking and protection from light. Next, three washes with TBST (10 min/wash) were performed. The membrane was then incubated with IRDye 680RD goat anti-rabbit (LI-COR) (1:20000, diluted in 1% nonfat dry milk-TBS) by incubating at RT for 3 h, protected from light.

Following the incubation with secondary Ab, the membrane is washed three times, 5 min each, with TBST then once with TBS for 5 min. Imaging was done with the LI-COR Odyssey imaging (Quality = medium; Resolution = 169 µm; Focus = 0 mm). The band density of each target was analyzed in FIJI ImageJ and normalized to total protein detected via Revert^TM^ 700 Total Protein Stain (**Fig. S4A**).

As additional normalization control for quantitation, blots probed with 4G10 as above were simultaneously probed with anti-β-actin monoclonal Ab from rabbit (Cell Signaling Technology) (1:7000, diluted in 1% nonfat dry milk-TBS). The β-actin bands were detected simultaneously with 4G10 (2-color fluorescence western blot) using IRDye680RD goat anti-rabbit (LI-COR) secondary antibodies (1:20000, diluted in 1% nonfat dry milk-TBS), by incubating at RT for 3 h. In addition to the normalization to Revert^TM^ 700 Total Protein Stain, the band density of target proteins was also normalized to the band density of β-actin (**Fig. S4B**). Results obtained from normalizing to Revert^TM^ 700 Total Protein Stain (Fig. 6B, 7C, 8, and 9B and 9C) were equivalent to the results obtained from normalizing to β-actin (data not shown).

To normalize tyrosine phosphorylated-Syk signal to total Syk, an equal amount of each sample, including the ladder, was duplicated on the same gel, proteins were separated via the SDS PAGE, and the gel was transferred onto nitrocellulose as described above. The membrane was cut into two halves (one for 4G10 and the other for total Syk); one half was processed via the above procedures described for probing with anti-phosphotyrosine Ab 4G10. The other half of the membrane was blocked with UltraCruz^®^ blocking reagent (Santa Cruz Biotechnology) for 1 h at RT with gentle shaking, with protection from light. Then it was washed three times, 5 min each, with 0.1% TBST. After washing, this half-blot was probed with anti-total Syk monoclonal Ab from mouse (1:100, diluted in UltraCruz^®^ blocking reagent, Santa Cruz Biotechnology) by incubating at RT for 1 h with gentle shaking. Following the incubation, the membrane was washed three times with 0.1% TBST and incubated with anti-mouse secondary Ab anti-IgG kappa binding protein (m-IgGκ BP) conjugated to CruzFluor™ 680 from goat (1:15000, diluted in UltraCruz blocking reagent, Santa Cruz Biotechnology) for 1 h at RT with shaking. The band density of phospho-Syk is normalized to the band density of total Syk (**Fig. 7B**); this result is equivalent in effect to the results obtained by normalizing the phospho-Syk to total protein (**Fig. 7C**). It was necessary to duplicate the samples on a single blot and cut the blot because the 4G10 Ab and the total Syk required different reagents and incubation conditions. Protein transfer onto the membrane was always efficient and consistent across all lane]s (**Fig. S4A**); thus, duplicating and cutting the blot is more effective than stripping and reprobing, which can result in signal loss^105,106^.

Bands were analyzed in FIJI ImageJ. Brightness but not contrast was adjusted as necessary. Protein bands to analyze were located based on molecular weight, and a region of interest was drawn both around the bands (across lanes) and around an equivalent area in a blank lane; mean fluorescence per pixel was measured, and the background was subtracted. Duplicates (on the same blot) were averaged, and data were normalized to Revert^TM^ 700 Total Protein Stain and then to Antigen-alone samples. Linearity tests were performed as required for quantitative western blotting ^104^ (**Fig. S5A-F**). The amount of protein loaded into each sample lane on the gel, 12 µg as noted above, was chosen as being within the linear range of all target proteins, including total protein signal (determined using Revert^TM^ 700 Total Protein Stain) with R-squared values ranging from 0.94 – 0.99.

### Syk Tyr phosphorylation ELISA

The PathScan^®^ Phospho-Syk (panTyr) sandwich ELISA (Cell Signaling Technology) was used to measure levels of phosphorylated Syk according to the manufacturer’s instructions. This pan-Tyr ELISA detects phosphorylation at all Syk’s Tyr residues. For this assay, RBL-2H3 cells were plated at cell density of 1.1375×10^7^ cells (in 6.5 mL RBL media) per 10 cm diameter Corning^®^ treated culture dishes (CellBIND^®^ surface, Corning) and were incubated overnight at 37 °C/5% CO_2_. The following day, cells were sensitized with anti-DNP mouse IgE (0.1 µg/mL) (Millipore Sigma) for 1 hour at 37 °C/5% CO_2_. Following sensitization, cells were washed twice with warm BT (5 mL/dish) and then pre-treated with 10 µM CPC (for samples labeled “CPC” on the graph) or BT (15 mL/dish) for 30 min at 37 °C/5% CO_2_. At the end of the pre-treatment, liquid was removed from the wells, and cell groups “Spont.,” “Ag,” and “Ag + CPC” were stimulated with 15 mL/dish pre-warmed BT, Ag (0.005 or 0.0007 µg/mL), or Ag (0.005 or 0.0007 µg/mL)/CPC, respectively, for 5 min at 37 °C. After this stimulation period, cell dishes were immediately inverted to remove the treatment, placed on ice, and washed with ice-cold PBS (5 mL/dish) with shaking, and the wash was thoroughly removed. Cells were lysed on ice for 5 min with 400 µL/dish ice-cold lysis buffer (Cell Signaling Technology) containing protease inhibitor phenylmethanesulfonyl fluoride (PMSF) at 1 mM (Millipore Sigma, CAS no. 329-98-6). Cells were harvested into prechilled, sterile 15 mL centrifuge tubes by thoroughly scraping the bottom of each dish with disposable sterile cell scrapers. Samples were mechanically lysed via probe sonication (Branson SFX 250 Sonifier) (Pulse, Time, On = 1 sec, Off = 3 sec, Total on = 3 sec, Amplitude: 50%). Sample lysates were transferred to pre-chilled, sterile microcentrifuge tubes and spun (Eppendorf, Centrifuge 5417R, rotor F-45-30-11) at 16,600 x g for 10 min at 4 °C, and the supernatant was transferred to new pre-chilled microcentrifuge tubes. Sample lysate supernatant (100 µL/sample) was transferred into each ELISA microwell (which was precoated with anti-Syk monoclonal Ab), and the plate was sealed with parafilm, wrapped in aluminium foil to protect from light, and incubated overnight (∼18 hours) at 4 °C. The following day, liquid was removed from wells, which were washed 4X with ice-cold assay wash buffer (200 µL/well per wash). Samples were then probed with a phospho-Syk mouse detection primary Ab, an HRP-linked secondary Ab, followed by substrate per manufacturer’s instructions. The absorbance was immediately measured in Corning Costar clear flat bottom plates at 450 nm by a plate reader (BioTek, Synergy 2). The average absorbance from the “Spont.” unstimulated group (representing background signal at 450 nm and constitutive phosphorylation) was subtracted from the absorbance readings of the stimulated groups (Ag, Ag + CPC). Finally, values were normalized to to Antigen-alone samples.

### Lyn Y507 ELISA

The RayBio^®^ human phospho-LYN (Y507) ELISA (Ray Biotech^®^) was used to measure the extent of phosphorylation of Lyn’s negative regulatory site, Y507, according to the manufacturer’s instructions. The procedure was the same as for the Syk ELISA detailed above, except that Ag was applied for a 2-min stimulation period, 1200 µL/dish ice-cold lysis buffer (with 1.2 mM PMSF) was used in place of 400 µL/dish, and (due to high signal) the sample lysate supernatants were diluted 1:200 in assay buffer before application to the ELISA. After subtraction of signal from a no-cell control well that was processed in tandem with all other samples, the phosphorylation signal for each group was normalized to Spont. control.

## Statistical analyses

All analyses were performed in Graphpad Prism. Mean ± SEM was determined by averaging biological replicates from at least three independent days of experiments. To determine significance, Student’s t-tests and ANOVA with Tukey’s post hoc tests were used.

## Results

### CPC inhibits degranulation in LAD2 human mast cells

Cells were sensitized with human IgE for 24 h, washed with BT, and pretreated with CPC in BT for 1 h (“1.5 h CPC”) or 30 min (“1 h CPC”). After CPC pretreatment, cells were stimulated with 2.5 ng/mL (**Fig. 2A**) or 10 ng/mL (**Fig. 2B**) streptavidin ± CPC for 30 min, and levels of degranulation were determined using a fluorescence-based assay as previously described^18^. The absolute degranulation values following stimulation with 2.5 (**Fig. 2A**) and 10 ng/mL (**Fig. 2B**) streptavidin are 28% ± 2% (SEM) and 32% ± 1% (SEM), respectively. The basal level of degranulation from unstimulated cells, or spontaneous (Spont.), is 9% ± 2% (SEM). The data show that CPC significantly inhibits the degranulation in LAD2 human mast cells following stimulation (**Fig. 2**), with somewhat stronger inhibition at the lower streptavidin dose (based on the assessment of statistical significance indicated by the number of asterisks). By 10 μM, CPC reduces degranulation levels down to Spont. levels (**Fig. 2A & 2B**). CPC inhibits degranulation in a dose-dependent manner (**Fig. 2**). CPC’s inhibition of LAD2 degranulation is also time-dependent, with the longer CPC exposure time of the “1.5 h CPC” group in **Fig. 2B** resulting in greater inhibition of degranulation than the “1 h CPC” group.

### CPC’s inhibition of mast cell degranulation is not specific to the type of crosslinker used to aggregate FcεRI IgE receptors

Previously, we showed that CPC inhibits RBL mast cell degranulation stimulated by multivalent Ag^17,18^. The current study probed the effects of CPC on degranulation when a bivalent crosslinker (anti-IgE IgG) is used in the place of Ag for the crosslinking of IgE-bound FcεRI receptors. Each anti-IgE antibody can bind to two IgE-bound FcεRI receptors simultaneously, while the multivalent Ag (DNP-BSA) can bind and aggregate more than two IgE-FcεRIs (**Fig. 3**).

**Fig 3.**
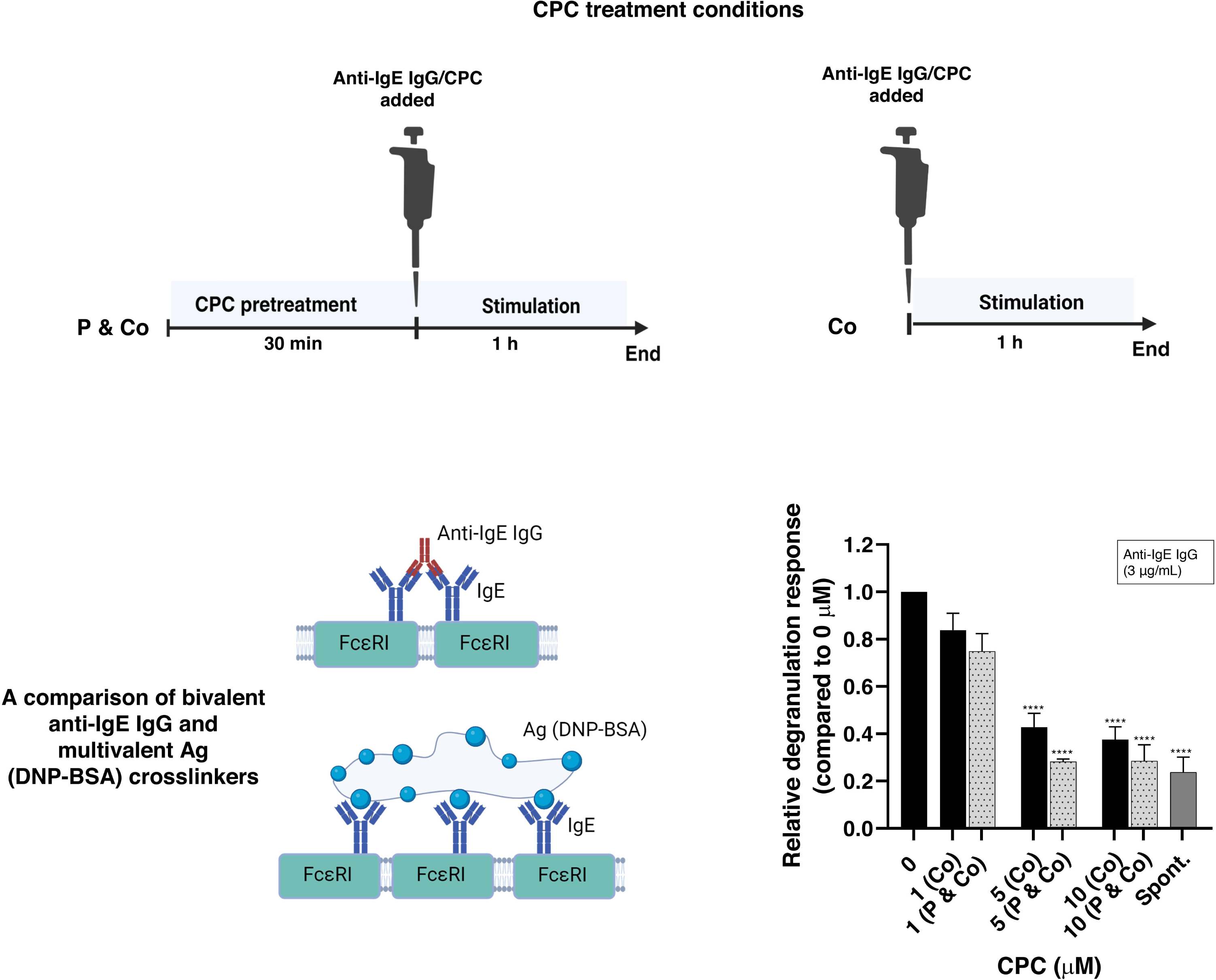
Degranulation responses of RBL mast cells stimulated by anti-IgE IgG crosslinking and exposed to micromolar doses of CPC. Cells were sensitized for 1 h with mouse IgE. For spontaneous release (“Spont.” on x-axis), cells that had not been sensitized with IgE were exposed to BT for 1 h without anti-IgE IgG. Samples denoted “P & Co” (grey bars; see treatment schematic) had CPC pretreatment for 30 min before the 1 h anti-IgE IgG/CPC co-exposure. Samples denoted “Co” (black bars; see treatment schematic) did not have CPC pretreatment ahead of the 1 h anti-IgE IgG/CPC co-exposure. Cells were stimulated with 3 μg/mL anti-IgE IgG for 1 h. Values presented are means normalized to the control, ± SEM, of at least 3 independent days of experiments; each experiment was performed in triplicate. Statistically significant results compared to the 0 μM control of each graph are represented by ****p < 0.0001, determined by one-way ANOVA followed by Tukey’s post-hoc test.

Cells were IgE-sensitized, washed with BT, and pretreated with CPC (“P & Co”) in BT for 30 min before being stimulated with anti-IgE IgG (3 µg/mL) ± CPC for 1 h. CPC groups that were not CPC-pretreated ahead of antibody stimulation are referred to as “Co.” Degranulation levels were determined using a fluorescence-based assay as previously described^18^. The anti-IgE IgG dose used in these experiments, 3 µg/mL, elicited an average absolute degranulation response of 10% ± 2% (SEM), which is the same level elicited by a low dose of Ag (0.0001 μg/mL)^18^. Regardless of crosslinker used to elicit 10% absolute degranulation (whether bivalent anti-IgE in **Fig. 3** or 0.0001 μg/mL multivalent DNP-BSA^18^, CPC (5 or 10 μM) inhibits degranulation down to levels equivalent to spontaneous release “Spont.” (according to ANOVA with Tukey’s post hoc test).

### CPC inhibits degranulation due to its pyridinium group’s cationic nitrogen

Hexadecylbenzene (HDB) is chemically structurally similar to CPC, apart from its positively-charged nitrogen in the head group ring^22^, as shown in **Fig. 4A**. To isolate the relevance of the quaternary nitrogen and positive charge of CPC, compared with the other structural features (saturated lipid tail and conjugated ring), we assessed HDB’s effect on degranulation in RBL cells (**Fig. 4C**). CPC effect on mast cell degranulation, under conditions equivalent to those used for the HDB, was performed in tandem. Cells were IgE-sensitized, washed with BT, and pre-treated with CPC or HDB (see schematic in **Fig. 4**) in BT for 30 min before stimulation for 1 h with multivalent Ag (0.0004 μg/mL) ± CPC/HDB (**Fig. 4**). Degranulation levels were measured using a fluorescence-based assay as previously described^100^. Spontaneous samples (unstimulated cells) showed absolute degranulation levels of 1.8% ± 0.2% (SEM). Ag (0.0004 μg/mL) elicited absolute degranulation of 22% ± 1% (SEM) of granules released. HDB does not affect mast cell degranulation (**Fig. 4C**), even following HDB pretreatment. In contrast, CPC dramatically inhibits degranulation in a time- and dose-dependent manner (**Fig. 4B**. Thus, CPC’s mechanism of action may be mainly due to its pyridinium group’s positively-charged nitrogen.

**Fig 4.**
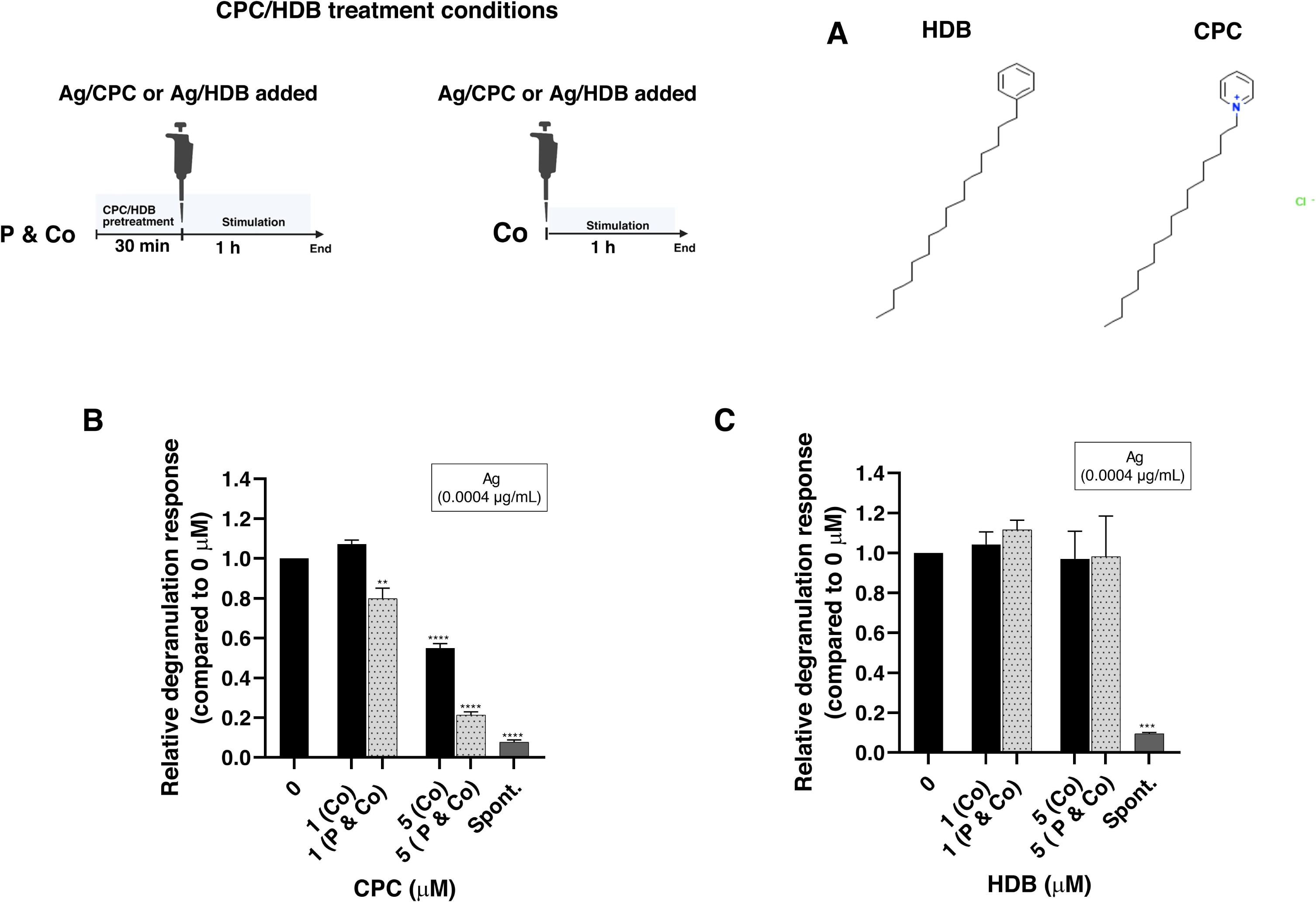
Relative degranulation response in RBL mast cells exposed to micromolar doses of CPC and HDB. **(A)** Chemical structures of HDB and CPC. Cells were sensitized with mouse IgE for 1 h. For spontaneous release (“Spont.” on the x-axis), non-IgE-sensitized cells were exposed to BT for 1 h without Ag. Samples denoted “P & Co” (grey bars; see treatment schematic) had CPC **(B)** or HDB **(C)** pretreatment for 30 min before the 1 h Ag/CPC or Ag/HDB co-exposure, respectively. Samples denoted “Co” (black bars; see treatment schematic) did not have CPC or HDB pretreatment ahead of the 1 h Ag/CPC or Ag/HDB co-exposure. Cells were stimulated with 0.0004 μg/mL Ag for 1 h. Values presented are means normalized to the control, ± SEM, of at least 3 independent days of experiments; each experiment was performed in triplicate. Compared to the 0 μM control, the statistically significant results are represented by **p < 0.01, ***p *<* 0.001, and ****p < 0.0001, determined by one-way ANOVA followed by Tukey’s post-hoc test.

### CPC inhibits global tyrosine phosphorylation in Ag-stimulated RBL mast cells

We investigated the effect of CPC on global tyrosine phosphorylation using the In-Cell Western (ICW) and Western Blotting (WB) assays (**Figs. 5 and 6**). Cells were sensitized with IgE, washed in BT, pretreated in CPC in BT for 30 min, and stimulated with Ag ± CPC for 5 min (see schematics in **Figs. 5 and 6**). In ICW, anti-phosphotyrosine antibody MultiMab was used to detect global tyrosine phosphorylation. Upon stimulation with Ag (1 µg/mL; a high dose used to maximize Tyr phosphorylation signal in ICW [dose-response data not shown]), there is a significant increase in global tyrosine phosphorylation (**Fig. 5**) over the unstimulated cells (Spont.) in ICW. CPC (10 µM) significantly decreases the Ag-stimulated tyrosine phosphorylation to constitutive levels equivalent to Spont. (according to ANOVA with Tukey’s post hoc test) (**Fig. 5**). As revealed by whole cell lysate Western Blot with anti-phosphotyrosine antibody 4G10, there is a high level of constitutive phosphorylation in unstimulated mast cells (**Fig. 6A**) and only a subset of proteins become Tyr phosphorylated upon Ag stimulation. CPC does not affect this high constitutive phosphorylation in mast cells (**Fig. S5**). Upon stimulation with Ag (0.0007 µg/mL (dose chosen for robust Tyr phosphorylation response and for CPC response^18^, there is a significant increase in global tyrosine phosphorylation (**Fig. 6A representative blot; Fig. 6B**, quantified entire lanes) over the unstimulated cells (Spont.) in WB. CPC (10 µM) significantly decreases the Ag-stimulated tyrosine phosphorylation (**Fig. 6B**).

**Fig 5.**
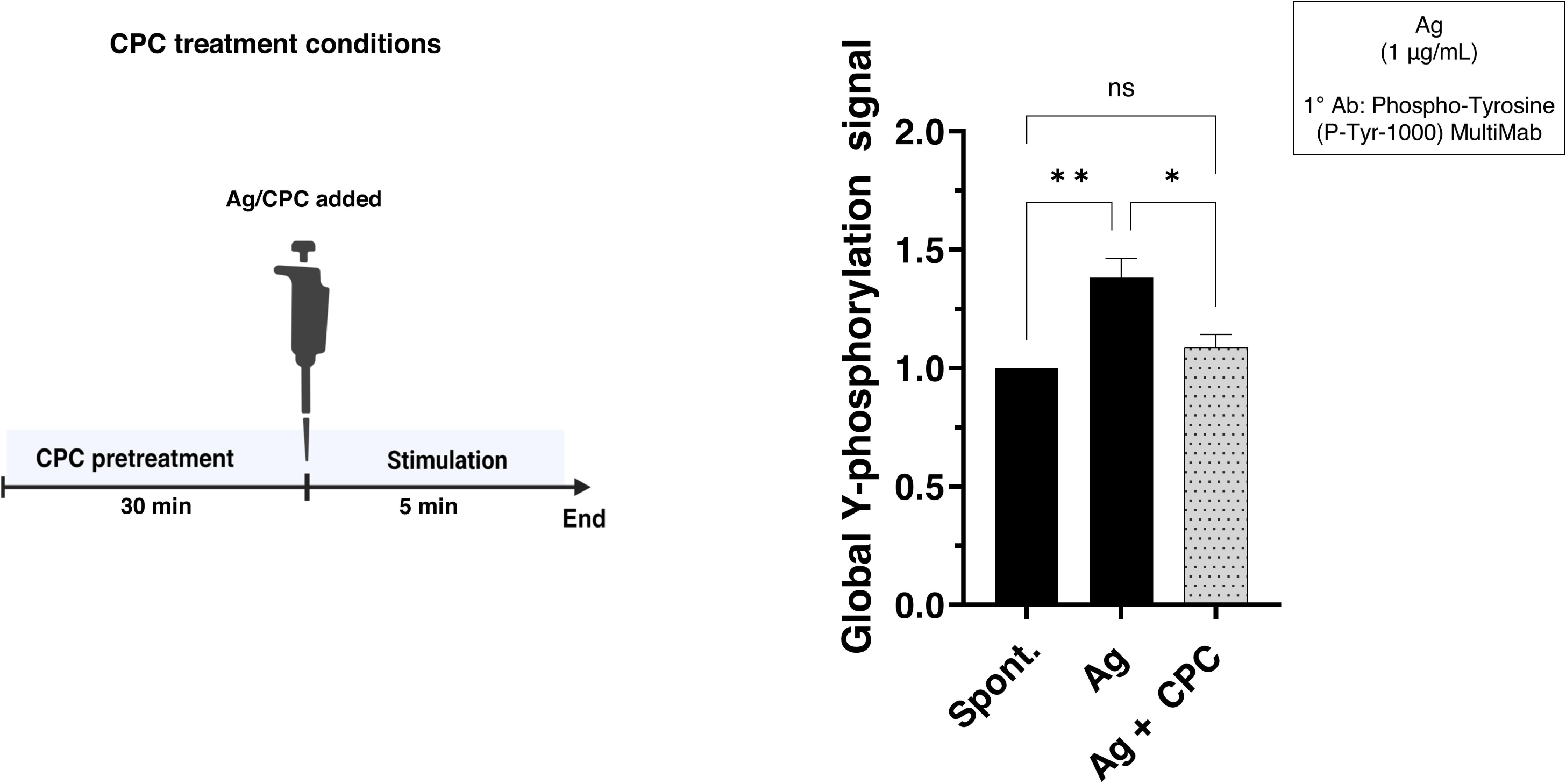
CPC effects on global tyrosine phosphorylation in Ag-stimulated RBL mast cells assessed via In-cell Western. Cells were sensitized with mouse IgE for 1 h. “Spont.” treatment represents non-IgE-sensitized cells that were exposed to BT for 5 min in the absence of Ag. Cells were pre-exposed to 10 µM CPC (or BT) for 30 min, followed by Ag (1 µg/mL) ± 10 µM CPC for 5 min at 37 °C. Tyrosine phosphorylation was quantified via ICW using a LI-COR Odyssey CLx: probing with the anti-phosphotyrosine MultiMab antibody, a fluorescent secondary antibody, and CellTag 700 to normalize cell number. Values presented are means normalized to Spont. group, ± SEM, of at least 3 independent days of experiments. Statistically significant results compared to the appropriate controls are represented by *p < 0.05 and **p < 0.01, determined by one-way ANOVA followed by Tukey’s post-hoc test.

### CPC inhibits tyrosine phosphorylation of Syk kinase

Syk is activated by Lyn kinase via tyrosine phosphorylation^23,59,107^, which is crucial to the degranulation pathway^62^. We probed the effect of CPC on Syk’s tyrosine phosphorylation in whole-cell lysates via ELISA (PathScan Phospho-Syk panTyr Sandwich ELISA kit, Cell Signaling Technology) and WB. Cells were sensitized with IgE, washed in BT, pretreated in CPC in BT for 30 min, and stimulated with Ag ± CPC for 5 min (see schematic in **Fig. 7**).

ELISA data shows that CPC inhibits the tyrosine phosphorylation of Syk elicited by moderate Ag (0.0007 µg/mL) stimulation (**Fig. 7A**) but does not significantly do so when a high Ag dose (0.005 µg/mL; see CPC degranulation effects and absolute degranulation percentage level elicited with this Ag dose^18^ is employed (**Fig. 7A**).

In WB, the anti-phosphotyrosine antibody 4G10 was used to detect tyrosine phosphorylation of Syk, and the tyrosine phosphorylated-Syk signal was normalized to total Syk (**Fig. 7B)** or to Revert Total Protein Stain (**Fig 7C**) in separate experiments. Upon stimulation with Ag (0.0007 µg/mL), there is robust stimulation (see representative blots in **Figs. 7B & C**) in tyrosine phosphorylation of Syk over the unstimulated group (“Spont.”) (2.9-fold in **Fig. 7B** and 2.4-fold in **Fig. 7C**). CPC (10 µM) inhibits the Ag-stimulated tyrosine phosphorylation of Syk (**Fig. 7B & C**), whether the tyrosine phosphorylated-Syk signal is normalized to total Syk (**Fig. 7B**) or normalized to Revert Total Protein Signal **(Fig. 7B).**

### CPC inhibits tyrosine phosphorylation of the adaptor molecule LAT

LAT is activated by Syk^108,109^ via tyrosine phosphorylation. We probed the effect of CPC on LAT’s tyrosine phosphorylation via WB using whole-cell lysates. Cells were sensitized with IgE, washed in BT, pretreated in CPC in BT for 30 min, and stimulated with Ag (0.0007 µg/mL) ± CPC for 5 min (see schematic in **Fig. 8**). The anti-phosphotyrosine antibody 4G10 was used to detect tyrosine phosphorylation of LAT based upon molecular weight. The tyrosine phosphorylated-LAT signal was normalized to Revert Total Protein Stain (**Fig. 8**). There is Ag stimulation (see representative blot in **Fig. 8**) in tyrosine phosphorylation of LAT over the unstimulated group (“Spont.”) (1.9-fold in **Fig. 8**). CPC (10 µM) inhibits the Ag-stimulated tyrosine phosphorylation of LAT **(Fig. 8)**.

### CPC does not alter the phosphorylation status of Lyn kinase

Lyn is activated via tyrosine phosphorylation^51,52^ (at Y397) and dephosphorylation^55,56^ (at Y507) upon crosslinking of the FcεRI receptors. We have investigated the effect of CPC on the tyrosine phosphorylation of Lyn (Y507) via ELISA and tyrosine phosphorylation of Lyn (total pY and Y397) in whole-cell lysates via WB. Cells were sensitized with IgE, washed in BT, pretreated in CPC in BT for 30 min, and stimulated with Ag ± CPC for 2 min (for Y507 ELISA **Fig. 9A** and Y397 WB **Fig. 9B**) or 5 min (for total pY WB **Fig. 9C**) (see schematic in **Fig. 9**).

CPC does not significantly affect Lyn phosphorylation at Y507 in whole-cell lysates as assessed via ELISA at two different Ag stimulation doses, though there is a hint of stimulation of this negative regulatory tyrosine by CPC (**Fig. 9A**).

Notably, there is no significant increase in phosphorylated-Lyn signal (Y397 and total Lyn, **Fig. 9B & C**, respectively) over the unstimulated group (Spont.) upon Ag stimulation. Thus, even in unstimulated whole mast cell lysates, Lyn exhibits a high level of constitutive tyrosine phosphorylation (see “no Ag” band in the representative blots beneath the graphs in **Fig. 9B** and **9C)**. CPC does not affect this level of phosphorylation, either at the positive regulatory Y397 or in the total phosphorylation signal across all of its tyrosines in these whole cell lysate samples (**Fig. 9B and 9C**).

## Discussion

CPC inhibits the functioning of immune cells^18,110^ despite being an agent tasked with aiding the human immune system to fight infection. In this study, we discovered an underlying mechanism of action of CPC in mammalian mast cells: it inhibits regulatory cascades of tyrosine phosphorylation.

Previously, we showed that CPC inhibits degranulation of rat RBL mast cells^17,18^. In this study, we confirm that CPC also inhibits degranulation of human LAD2 mast cells **(Fig. 2)**. Thus, CPC’s inhibition of degranulation is not exclusive to a particular type of mast cell and is applicable to human cells. CPC inhibits LAD2 degranulation in a time- and dose-dependent manner **(Fig. 2)**, as seen in RBLs^18^. Future experiments will assess the effect of CPC on tyrosine phosphorylation and Ca^2+^ mobilization in LAD2 human mast cells.

CPC’s inhibition of mast cell degranulation is not specific to the type of crosslinker used to aggregate FcεRI IgE receptors. As previously shown, CPC inhibits degranulation induced by multivalent Ag which crosslinks multiple receptors together^17,18^ (see schematic in **Fig. 3**), and as such, in this study, we show that CPC equivalently inhibits degranulation induced by a bivalent anti-IgE IgG (**Fig. 3**). It is known that CPC is a more potent inhibitor when lower crosslinker concentrations (which elicit lower amount of absolute degranulation) are used^18^, so it was important to compare anti-IgE and DNP-BSA results at equivalent amounts of stimulated degranulation in the control. Thus, the dose of anti-IgE IgG chosen for use in this in this study (**Fig. 3)** was chosen to match the cell stimulation potential of multivalent Ag DNP-BSA (0.0001 μg/mL) used in a previous study^18^: each elicited an average absolute degranulation response of 10% (**Fig. 3** and Obeng et al.^18^). Regardless of crosslinker used to elicit 10% absolute degranulation (whether bivalent anti-IgE in **Fig. 3** or 0.0001 μg/mL multivalent DNP-BSA^18^, CPC (5 or 10 μM) inhibits degranulation down to levels equivalent to spontaneous release “Spont.” (according to ANOVA with Tukey’s post hoc test). Therefore, CPC’s inhibition is not exclusive to a particular crosslinker. CPC inhibition of either anti-IgE IgG-induced or multivalent Ag-induced degranulation shows that CPC does not act via interference in the binding of a crosslinker to IgE and suggests that CPC’s inhibitory effect is independent of signaling platform receptor architecture (numbers and orientation of receptors aggregated). Rather, CPC interferes with a signaling component farther into the pathway leading to degranulation. In contrast, other quaternary ammonium compounds like benzalkonium chloride are known to inhibit MC degranulation by direct interference with cationic stimulants (polymers and peptide Substance P) at the cell surface; these quats do not inhibit Ag-stimulated degranulation^111^.

We investigated the mechanistic relevance of the quaternary nitrogen in the pyridinium headgroup of CPC^1^ (Fig. 4A) by comparing the effects of CPC to those of HDB on degranulation (**Fig. 4B & 4C**). CPC as low as 1 µM inhibits degranulation but HDB does not (**Fig. 4 B & 4C**). HDB is chemically similar to CPC but has a benzyl group instead of the quaternary nitrogen^22^; thus, CPC’s mechanism of action in mast cells depends on its positively-charged quaternary nitrogen in the pyridinium group.

The concentrations of HDB reported on the x-axis of **Fig. 4C** were determined using UV-Visible spectrophotometry and the extinction coefficient of CPC (4389 M^−1^ cm^-1^), as described in Materials and Methods. A published UV-Vis spectrum of HDB^96^ depicts a peak ∼260 nm corresponding to ∼251 M^−1^ cm^-1^ (if units are standard), but solvent used and units are not clearly indicated at the site. If 251 M^−1^ cm^-1^ or a similar, lower extinction coefficient is used in place of 4389 M^−1^ cm^-1^, in the calculations, the HDB concentration would be determined to be close to the nominal 106.6 µM (see Materials and Methods) instead of the reported concentration of ∼ 10 µM. Thus, it’s possible that the doses indicated on the x-axis of **Fig. 4C** are ∼10-fold lower than the actual doses received by the cells: it may be that ∼10 and 50 µM of dissolved HDB were tested rather than the reported 1 and 5 µM. In contrast, as noted in Materials and Methods, the reported CPC doses used in **Fig. 4B**, which are based on UV-Vis results, are very close to the nominal values (from grams dissolved into mL of liquid). Thus, even at high concentrations, HDB exhibits no effect on degranulation, and the CPC mechanism of action is due to the quaternary nitrogen in the pyridinium headgroup **(Fig. 4)**. Future experiments will investigate the contribution of the lipophilic tail (**Fig. 4A)** to CPC’s mechanism of action by assessing the effect of isolated pyridinium, without the lipophilic tail, on mast cell degranulation.

Due to our previous study showing CPC inhibition of Ca^2+^ mobilization^18^ as well as the HDB results suggesting an electrostatic mode of action, we hypothesized that CPC may act via interference with Ag-stimulated phosphorylation that is required upstream of IP_3_-driving ER Ca^2+^ mobilization. Thus, we investigated the effect of CPC on global tyrosine phosphorylation in Ag-stimulated mast cells via ICW (**Fig. 5)** and WB **(Fig. 6)** using whole-cell lysates. In both ICW (**Fig. 5**) and WB (**Fig. 6B**), there was a significant increase in the tyrosine phosphorylation signal of the Ag-stimulated group over the unstimulated group (Spont.) as expected.

**Fig 6.**
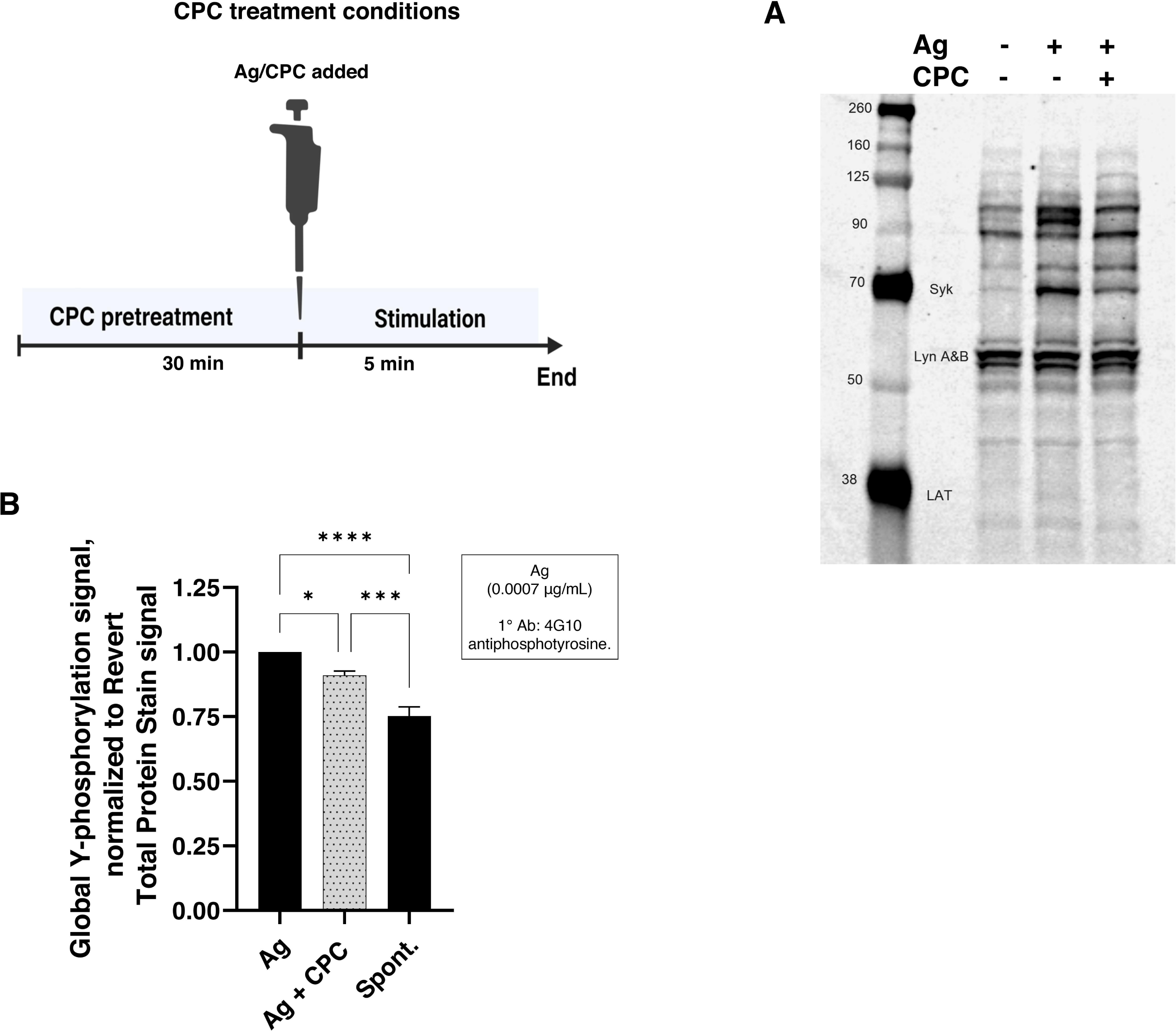
CPC effects on global tyrosine phosphorylation in Ag-stimulated RBL mast cells assessed via Western blot. Cells were sensitized with mouse IgE for 1 h. “Spont.” treatment represents cells that were exposed to BT for 5 min in the absence of Ag. Cells were pre-exposed to 10 µM CPC (or BT) for 30 min, followed by Ag (0.0007 µg/mL) ± 10 µM CPC for 5 min at 37 °C. For tyrosine phosphorylation detection via western blot, proteins in whole-cell lysate were separated via SDS-PAGE, transferred onto a nitrocellulose membrane overnight at 4 °C, blocked with 5% nonfat milk, probed with anti-phosphotyrosine antibody 4G10 overnight at 4 °C, and detected with IRDye 800CW goat anti-mouse secondary antibody. A representative western blot image of tyrosine-phosphorylated proteins in whole-cell lysate **(A)**. The density of the entire lane was quantified, and the Revert Total Protein Stain signal was used for normalization of quantitative data **(B)**. Values presented are means normalized to the Ag group, ± SEM, of at least 3 independent days of experiments. Statistically significant results compared to the appropriate controls are represented by *p < 0.05, ***p < 0.001, and ****< 0.0001 determined by one-way ANOVA followed by Tukey’s post-hoc test.

In the ICW (**Fig. 5**), a high Ag dose (1 µg/mL) was used in order to elicit significant Tyr phosphorylation signal over the unstimulated group^112^. Preliminary experiments revealed the efficacy of the anti-phosphotyrosine primary antibody MultiMab for ICW (4G10 did not work for ICW^112^). Upon stimulation with Ag, as revealed via WB (**Fig. 6A**), specific bands such as Syk (**Fig. 6A, 7B, and 7C**) and LAT (**Fig. 6A and 8**) are stimulated. Lyn displays high constitutive tyrosine phosphorylation (**Fig. 6A**) as expected^41,54^. While we have probed in detail the CPC effects on the earliest Tyr phosphorylation events (Lyn, Syk, LAT), we recognize that there are additional bands in the whole cell lysate that not only undergo Ag stimulation but that also are inhibited by CPC (**Fig. 6A**); these are under investigation. (Also, the Tyr phosphorylation of FcεRI itself was not analyzed because our electrophoresis blotting conditions, required to obtain a wide view of CPC effects on whole-cell Tyr phosphorylation, necessitated optimized electrophoresis and transfer conditions for proteins in the mid-range of molecular weight, rather than for the small subunits of FcεRIβ and FcεRIγ.)

**Fig 7.**
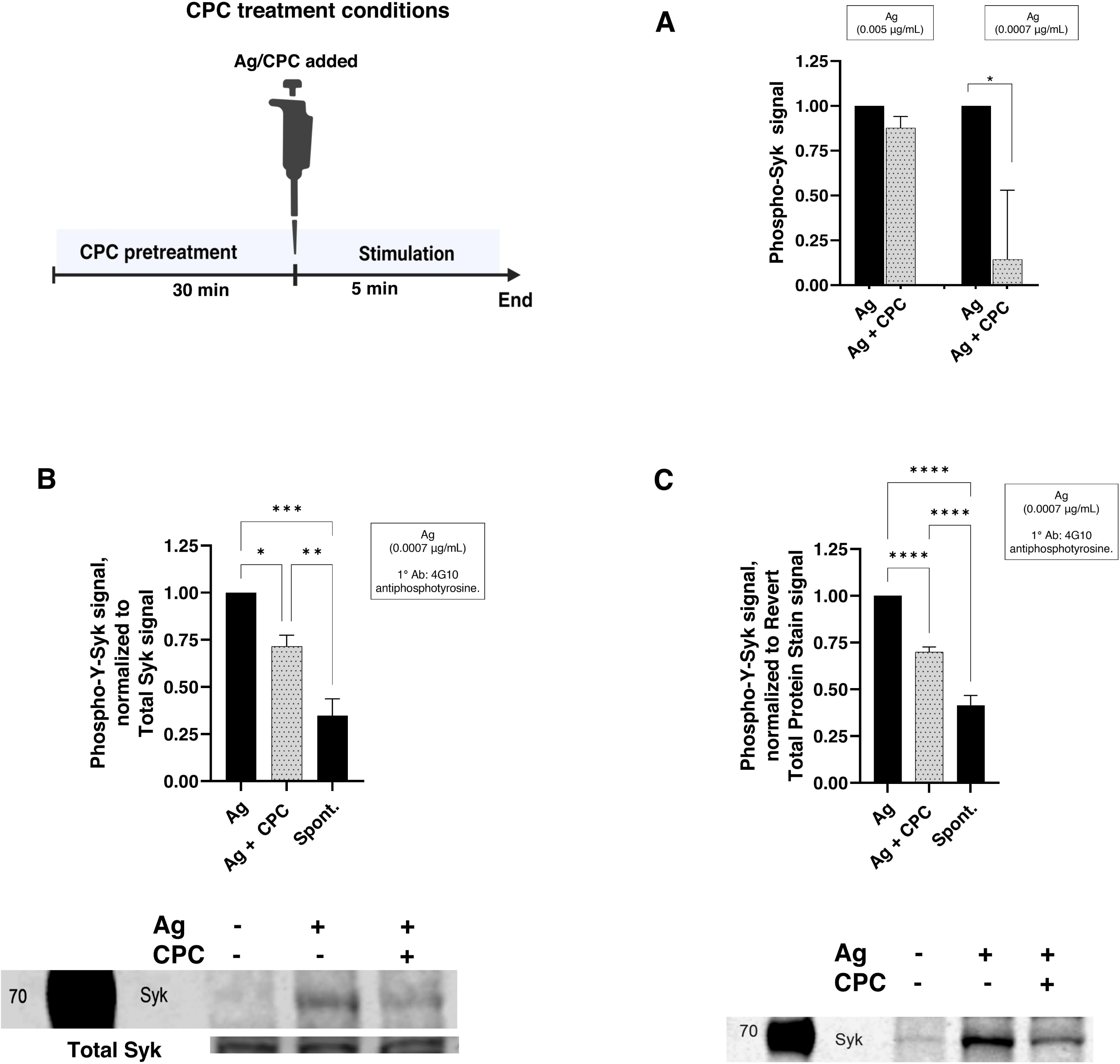
CPC effects on Syk kinase tyrosine phosphorylation in RBL mast cells. Cells were sensitized with mouse IgE for 1h. “Spont.” treatment represents cells that were exposed to BT for 5 min in the absence of Ag. Cells were pre-exposed to 10 µM CPC (or BT) for 30 min, followed by Ag (0.0007 or 0.005 µg/mL) ± 10 µM CPC for 5 min at 37 °C. PathScan phospho-Syk sandwich ELISA was used to measure Syk phosphorylation levels **(A).** For Syk tyrosine phosphorylation detection via western blot **(B)** & **(C)**, proteins in whole-cell lysate were separated via SDS-PAGE, transferred onto a nitrocellulose membrane overnight at 4 °C, blocked with 5% nonfat milk, incubated with anti-phosphotyrosine antibody 4G10 overnight at 4 °C, detected with IRDye 800CW goat anti-mouse secondary antibody. In **(B)**, total Syk antibody was used for normalization. To quantify the levels of total Syk, transferred proteins were blocked with UltraCruz blocking reagent, probed with an anti-Syk antibody at room temperature for 1 h, and detected with a goat anti-mouse secondary antibody conjugated to CruzFluor 680. In **(C),** Revert Total Protein Stain signal was used for normalization. Values presented are means normalized to the Ag group, ± SEM, of at least 3 independent days of experiments. Statistically significant results compared to the appropriate controls are represented by *p < 0.05, **p < 0.01, ***p < 0.001, and ****p < 0.0001, determined by one-way ANOVA followed by Tukey’s post-hoc test. Representative blot samples are shown below their corresponding bar graphs; numbers refer to molecular weight standard values.

Syk, ∼72 KDa, is crucial for degranulation^61,62^: activated Syk activates LAT and PLCγ1^63^, necessary for Ca^2+^ mobilization. In the absence of stimulation, Syk has no or little tyrosine phosphorylation^113^. Upon Ag stimulation, there is a robust increase in Syk tyrosine phosphorylation signal over the unstimulated group detected by WB (2.9-fold in **Fig. 7B** and 2.4-fold in **Fig. 7C**). CPC dramatically inhibits Ag-stimulated Syk phosphorylation (**Fig. 7B & C**). Syk’s tyrosine phosphorylation inhibition is further corroborated by CPC inhibition of Ag-stimulated Syk phosphorylation as detected with ELISA (**Fig. 7A**).

Upon Ag stimulation, six key tyrosine residues are phosphorylated in Syk: one found between the two SH2 domains (Y130), three in the linker domain (Y317, Y342, Y346), and two in the activation loop of the catalytic kinase domain (Y519 and Y520)^114^. The amino acid numbers given are for mouse, rat, chicken; the corresponding amino acids in human are +6 (e.g., Y519 in mouse = Y525 in human). The ELISA and WB methods used in the current study to assess Syk employed antibodies that detect all phosphorylated tyrosines on the kinase (see Materials and Methods).

Lyn catalyzes the phosphorylation of the three linker-domain tyrosines^114^, including Y317, which is a negative regulatory tyrosine^114^. While Lyn does not phosphorylate the catalytic-domain tyrosines 519/520, its phosphorylation of Syk’s tyrosines Y342 and Y346 is stimulatory. Indeed, Y342 and Y346 have been identified as docking sites for SH2 domain-containing molecules and are important for the binding of Syk to PLCγ1^115^ and thereby for the critical next step of PIP_2_-mediated Ca^2+^ mobilization. Syk autophosphorylates its Y130^113^ and Y519/Y520 in the kinase region. Phosphorylation of Y519/Y520 residues within Syk is critical for its ability to transmit downstream signals and lead to degranulation^107^. It is not yet known which Tyr within Syk are unphosphorylated due to the presence of CPC, but the majority of Syk’s phosphorylated tyrosines are positive-regulatory residues, and much of the phosphorylation signal is abrogated by CPC (**Fig. 7**). Thus, future experiments will pinpoint the key affected residues by specifically testing CPC effects on Ag-stimulated Y519/Y520 phosphorylation with antibodies specific to those residues (ELISA and WB). Also, future experiments will test whether HDB affects Syk phosphorylation.

Once Syk is inhibited by CPC, downstream events such as LAT activation are affected, in line with the previously-observed suppression of ER Ca^2+^ efflux, cytosolic Ca^2+^ rise, and, subsequently, degranulation^18^. LAT is activated by Syk via tyrosine phosphorylation^63,109^. LAT tyrosine phosphorylation is increased upon Ag stimulation, compared to the unstimulated group (Spont.), 1.9-fold increase (**Fig. 8**). CPC significantly inhibits the Ag-stimulated tyrosine phosphorylation of LAT (**Fig. 8**), a result expected as a natural consequence of CPC’s inhibition of Syk, LAT’s activating enzyme. Syk-activated LAT is required to activate PLCγ1^63^, needed for IP_3_ generation and Ca^2+^ mobilization. Thus, CPC inhibition of LAT is a contributor to CPC’s suppression of Ca^2+^ mobilization and, downstream, degranulation^18^.

**Fig 8.**
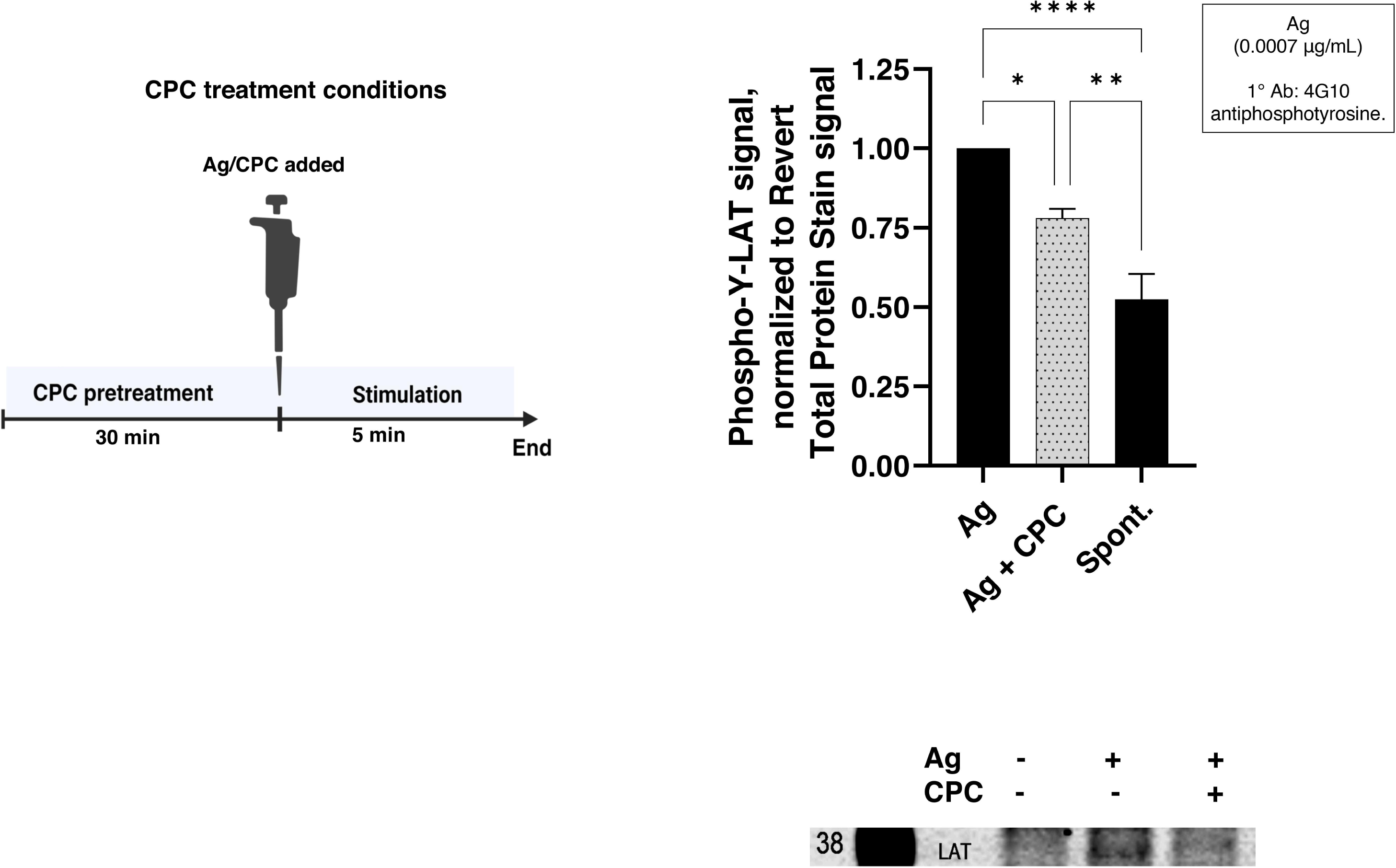
CPC effects on adaptor protein LAT tyrosine phosphorylation in RBL mast cells. Cells were sensitized with mouse IgE for 1h. “Spont.” treatment represents cells that were exposed to BT for 5 min in the absence of Ag. Cells were pre-exposed to 10 µM CPC (or BT) for 30 min, followed by Ag (0.0007 µg/mL) ± 10 µM CPC for 5 min at 37 °C. For LAT tyrosine phosphorylation detection via western blot, proteins in whole-cell lysate were separated via SDS-PAGE, transferred onto a nitrocellulose membrane overnight at 4 °C, blocked with 5% nonfat milk, incubated with anti-phosphotyrosine antibody 4G10 overnight at 4 °C, and detected with IRDye 800CW goat anti-mouse secondary antibody. Revert Total Protein Stain signal was used for normalization. Values presented are means normalized to the Ag group, ± SEM, of at least 3 independent days of experiments. Statistically significant results compared to the appropriate controls are represented by *p < 0.05, **p < 0.01, and ****p < 0.0001, determined by one-way ANOVA followed by Tukey’s post-hoc test. Representative blot samples are shown below their corresponding bar graphs; numbers refer to molecular weight standard values.

Lyn is upstream of Syk: upon Ag crosslinking of IgE-FcεRI, immunoreceptor tyrosine-based activation motifs (ITAMs) in the cytoplasmic tails of the high-affinity IgE receptor FcεRI’s β and γ subunits^39,40^ become phosphorylated by the tyrosine kinase Lyn^40,41^. Once phosphorylated, the negatively-charged ITAMs recruit Syk via its positively-charged SH2 domains^42,44,45,58,59^.Lyn then activates Syk^58,59^. We examined CPC’s effect on tyrosine phosphorylation of Lyn in whole-cell lysates, via ELISA (**Fig. 9A)** and WB **(Fig. 9B & C).**

CPC does not affect the tyrosine phosphorylation level of total Lyn in whole cell lysate samples under the conditions tested (**Fig. 9**). Lyn is known to possess relatively high constitutive Tyr phosphorylation^41,54,116^, a result we have confirmed here (**Figs. 6A, 9**). There are seven tyrosines in Lyn annotated as phosphorylated (ncbi.nlm.nih.gov/protein). While two of these are well-characterized for their roles in Lyn activation (positive-regulatory Y397, the phosphorylation of which drastically increases the activity of Lyn^117^; and negative-regulatory Y507, the phosphorylation of which decreases the activity of Lyn^118^, the large overall number of phosphorylated tyrosines, which may not change significantly due to Ag stimulation, could mask changes in Y397 and Y507 phosphorylation levels when viewed with a pan-pY antibody (4G10) from whole cell lysates (as in **Fig. 6A**). Moreover, in the case of the total Lyn’s Tyr phosphorylation (**Fig. 6A and 9C**) at a particular time point, the signal due to Ag-stimulated increase in phosphorylation of Y397^50–52^ may be neutralized by the Ag-stimulated decrease in tyrosine phosphorylation signal due to dephosphorylation of Y507^55,56^. Thus, in order to more clearly probe CPC effects on the positive- and negative-regulatory tyrosines, further experimentation was performed. We carried out WB with an anti-phosphoY397 Lyn antibody (**Fig. 9B**) but still did not register Ag stimulation or CPC disruption of the phosphorylation signal at this specific, activating residue. Nevertheless, ELISA results specific to Y507 of Lyn suggest a hint of CPC stimulation of Y507 phosphorylation–which could mean CPC stimulation of the negative regulation of Lyn. While multiple timepoint experiments were conducted to optimize timing conditions (data not shown) and the 2 min timepoint for assessing Lyn Y507 (**Fig. 9A)** Y397 (**Fig. 9B**) was chosen based on published data from another cell type^55^, it is possible that we missed the optimal time for detecting Ag and/or CPC modulation of the Tyr phosphorylation levels of Lyn Y397 and Y507 in these cells.

**Fig 9.**
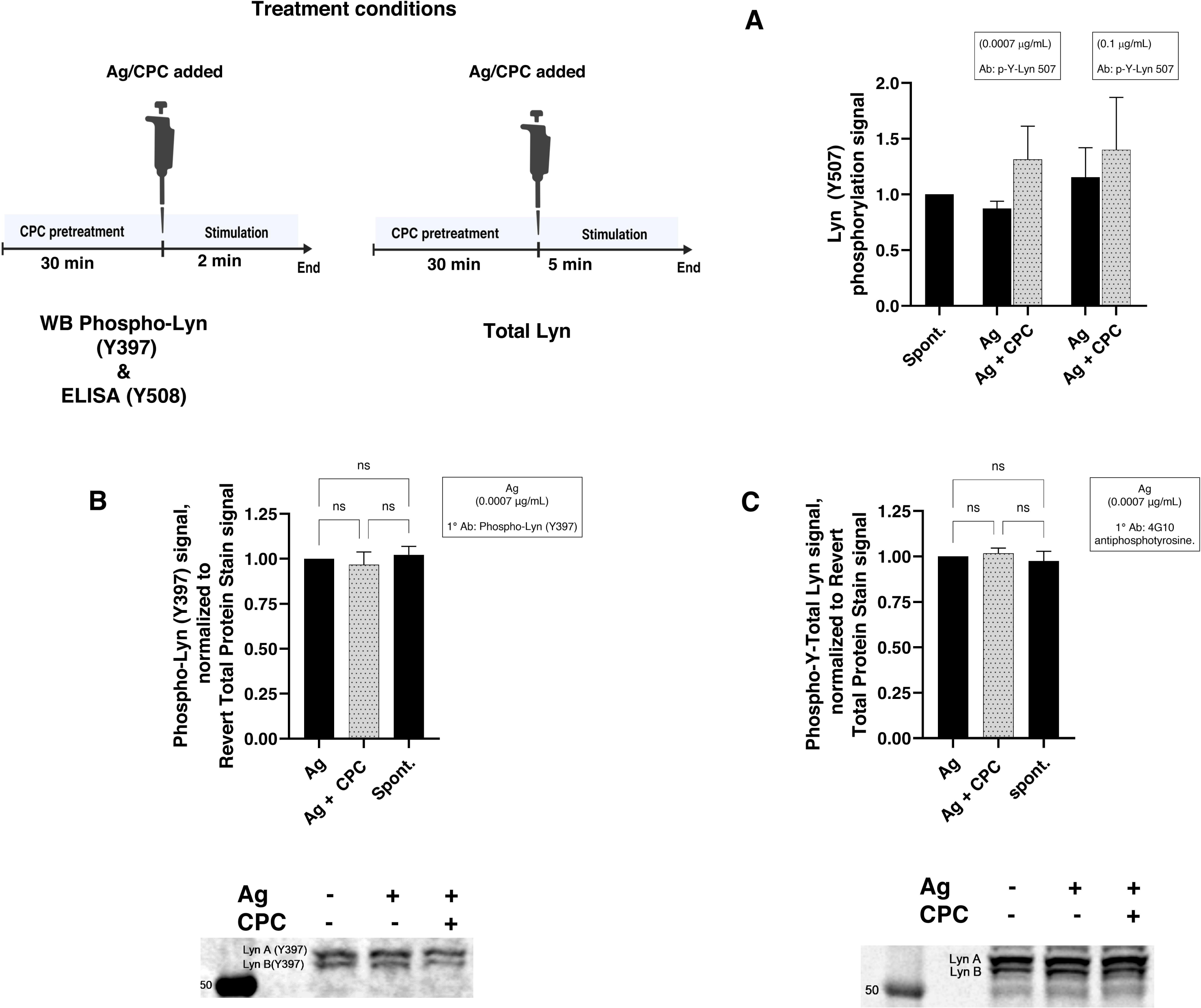
CPC effects on Lyn kinase tyrosine phosphorylation in RBL mast cells. Cells were sensitized with mouse IgE for 1 h. “Spont.” treatment represents cells that were exposed to BT in the absence of Ag. Cells were pre-exposed to 10 µM CPC (or BT) for 30 min, followed by Ag (dose as indicated on graphs) ± 10 µM CPC for 2 min **(A)** and **(B)** or 5 min **(C)** at 37 °C. Ray Biotech phospho-Y507 Lyn sandwich ELISA was used to measure Lyn Y507 phosphorylation levels **(A).** For Lyn tyrosine phosphorylation detection via western blot, proteins in whole-cell lysate were separated via SDS-PAGE, transferred onto a nitrocellulose membrane overnight at 4 °C, blocked with 5% nonfat milk, incubated with phospho-Lyn (Y397) antibody **(B)** or anti-phosphotyrosine antibody 4G10 **(C)** overnight at 4 °C, and detected with IRDye 700CW goat anti-rabbit and IRDye 800CW goat anti-mouse secondary antibodies, respectively. Revert Total Protein Stain signal was used for normalization. Values presented are means normalized to the Ag group, ± SEM, of at least 3 independent days of experiments. One-way ANOVA followed by Tukey’s post-hoc test revealed no statistically significant differences. Representative blot samples are shown below their corresponding bar graphs; numbers refer to molecular weight standard values.

Another possible explanation for the lack of apparent CPC effects on Lyn phosphorylation is that the preparation of homogenized whole cell lysates perhaps obscures the transitory Ag-induced changes in Lyn phosphorylation which likely occur in specific liquid-ordered (Lo) microdomains of the plasma membrane ^53,119^ which may represent only a small percentage of plasma membrane area and of Lyn molecules in the cell. Lyn isolated from these microdomains, which include crosslinked FcεRI receptors and exclude transmembrane phosphatases, possesses enhanced activity which is correlated with increased phosphorylation of the activating Y397^53^. The whole cell lysis likely scrambles the pools of Lyn and disallows precise analysis of these phosphorylation sites upon Ag stimulation and CPC treatment. Future efforts will focus on Lyn Y397 phosphorylation within these microdomains to better pinpoint CPC’s effect on Lyn.

A recent report using live-cell super-resolution microscopy showed a clear correlation between fluorescently-tagged probes’ preference for liquid-ordered (Lo) regions of membrane vesicles and their enrichment in B cell receptor clusters within live B cells^120^, lending strong evidence for the existence of “lipid rafts” or Lo membrane domains in living cells and corroborating their immune cell signaling significance. They used fluorescently-tagged probes such as one bearing the palmitate and myristoyl anchors of Lyn kinase (which, with saturated fatty acid chains, exhibits preference for Lo regions) (“PM probe”) in order to probe their position in the plasma membrane and their connection to immune cell signaling platforms while simultaneously knowing their tendency to cluster in Lo regions of model membranes. In fact, the authors showed that 16-C saturated chain hexadecanol stabilizes Lo-correlated microdomains in live B cells and thereby enhances phosphorylation of the B cell receptor, with similar effects observed in T cells as well^120^. CPC possesses the same 16-C saturated fatty acid chain as does hexadecanol, perhaps enabling it also, like hexadecanol, to interact with microdomains of clustered IgE receptors, where it may then disrupt FcεRI/Lyn/Syk interactions due to its positively-charged pyridinium group, leading to the Syk inhibition we have observed (**Fig. 7**). Lyn kinase (with its 16C membrane anchor), the PM probe used, the hexadecanol chemical modifier, and CPC each possess the same 16 saturated lipid chain. The work of Shelby *et al.* combined with our findings suggests that CPC might be preferentially intercalated into Lo “lipid raft” regions of clustered FcεRI and thereby able to exactly interfere with these precise signaling platform processes in mast cells. Please note, though, there are published instances of Lyn phosphorylation (at Y397) measured from whole lysate samples being inhibited by a drug, for example Luxeptinib^116^ despite the homogenized nature of those samples like ours. Thus, it remains a distinct possibility that CPC simply does not affect Lyn phosphorylation.

Future planned experiments will examine CPC effects on function of T cells, which share very similar upstream signaling machinery with mast cells. T cell signaling^121,122^ begins with T-cell receptor (TCR) aggregation^123,124^ analogous to FcεRI activation in MC. Tyrosine kinase (Lck) is analogous to Lyn in MC, bound to the PM and containing an SH2 domain to bind TCR ITAMs^125^. TCR activation involves tyrosine phosphorylation of Lck, Zap70, LAT^126^, leading to PLCγ activation, IP_3_ generation, and SOCE^127,128^, as in MC. PIP_2_ signaling is crucially important in T cells^129^. Thus, we hypothesize that CPC will similarly inhibit T cell function as it does MC degranulation.

Ag-stimulated tyrosine phosphorylation events culminate in the activation of PLCγ, which generates IP_3_ from PIP_2_. IP_3_ generation then triggers ER Ca^2+^ efflux, which is necessary for degranulation and which we earlier showed is inhibited by CPC^18^. Thus, future experiments will examine CPC effects on the activation of PLCγ and on the generation of IP_3_ in activated mast cells.

In conclusion, this study shows that CPC inhibits mast cell function, at doses ∼100X lower than found in consumer products, by disrupting specific tyrosine phosphorylation events. Here, we have extended the rodent MC findings to human MCs (LAD2). Previously we showed that CPC inhibits Ca^2+^ mobilization^18^, leading to the suppression of microtubule polymerization^18^ and, ultimately, degranulation^18^. CPC’s inhibition of Ca^2+^ mobilization and degranulation is independent of signaling platform receptor architecture (bivalent vs. multivalent crosslinking). We delineate the underlying biochemical mechanisms leading to the CPC’s repression of Ca^2+^ mobilization and degranulation in Ag-stimulated MCs. CPC inhibits degranulation due to its positively-charged quaternary nitrogen in the pyridinium headgroup (**Fig. 10**). CPC electrostatically disrupts and interferes with tyrosine phosphorylation events, particularly Syk and LAT (**Fig. 10**), which precede Ca^2+^ mobilization in the pathway. Thus, CPC thwarting tyrosine phosphorylation of Syk and LAT accounts for its inhibition of Ca^2+^ mobilization and degranulation (**Fig. 10**). CPC is a signaling toxicant, and any cell that employs a tyrosine phosphorylation cascade may be susceptible to its toxicity.

**Fig. 10.**
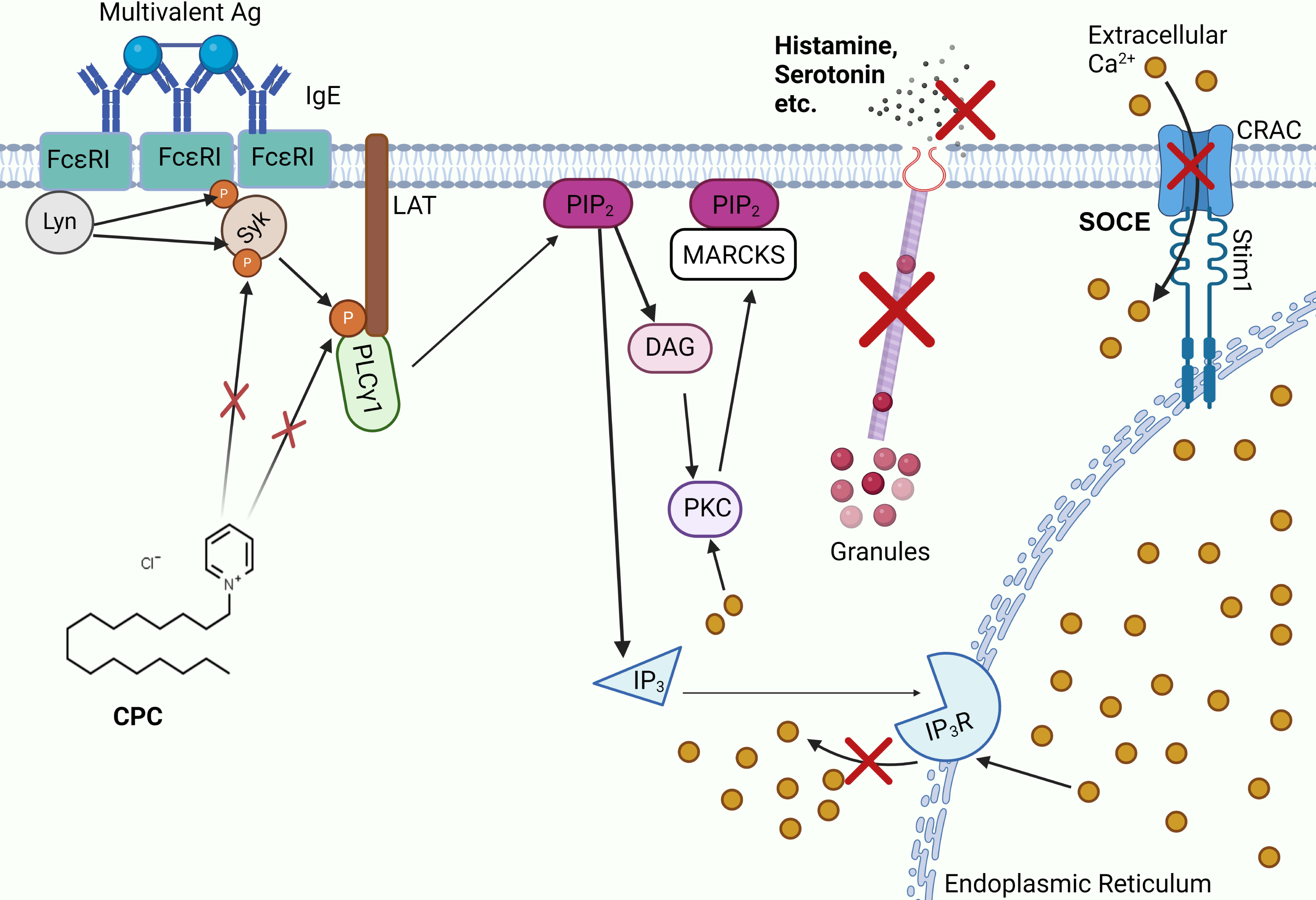
Summary of CPC targets in the mast cell signaling pathway. Following crosslinking of IgE-bound FcεRI receptors, tyrosine kinase Lyn phosphorylates FcεRI. Phosphorylated FcεRI recruits another Src family kinase, Syk, which is then phosphorylated by Lyn. Syk is an activator of adaptor hub LAT, which serves as a scaffold for activation of PLCγ1, which is essential for Ca^2+^ mobilization. In healthy mast cells, these early tyrosine phosphorylation events are necessary and precede Ca^2+^mobilization, a key mediator of the degranulation pathway. However, in this manuscript we have shown that CPC inhibits tyrosine phosphorylation of Syk and LAT. In a previous article, we showed that CPC suppresses Ca^2+^ mobilization out of the ER and through the plasma membrane CRAC channel into the cytosol, as well as the Ca^2+^-dependent downstream events microtubule polymerization and degranulation. The details of CPC effects between LAT and Ca^2+^ efflux from the ER remain to be elucidated. Created via BioRender.

## Abbreviations

Ag: antigen
BSA: bovine serum albumin
BT: Tyrodes-bovine serum albumin
CMC: critical micelle concentration
CPC: cetylpyridinium chloride
DAG: diacylglycerol
DMSO: dimethyl sulfoxide
DNP: dinitrophenyl
ER: endoplasmic reticulum
HDB: hexadecyl benzene
ICW: In-Cell Western
IgE: immunoglobulin E
IgG: immunoglobulin G
IP_3_: inositol 1,4,5-triphosphate
ITAMs: immunoreceptor tyrosine-based activation motifs
LAD2: laboratory of allergic diseases
LAT: linker for activation of T cells, LDH, lactate dehydrogenase
MC: mast cell
PIP_2_: phosphatidylinositol 4,5-bisphosphate
PLCγ: phospholipase C gamma
PM: plasma membrane
RBL-2H3: rat basophilic leukemia cells, clone 2H3
SEM: standard error of the mean
SARS-CoV-2: severe acute respiratory syndrome coronavirus 2
SH2: Src-homology-2 domain
SH3: Src-homology-3 domain
SOCE: store-operated Ca^2+^ entry
Syk: spleen tyrosine kinase
TCR: T-cell receptor

## Acknowledgments

This work was supported by the National Institutes of Health (NIH) under grant number 1-R15-ES034567-01 (PI: Gosse) and under the Institutional Development Award (IDeA) P20GM0103423 (PI: Coffman). A Faculty United for Toxicology Undergraduate Research and Education (FUTURE) Committee faculty research grant, from the Society of Toxicology, also provided student research funding. University of Maine funding that supported this work included an Institute of Medicine (IoM) Summer Graduate Student Fellowship, an IoM Seed Grant, University of Maine System Research Reinvestment Fund Grant Program Track 1 Rural Health and Wellbeing Grand Challenge, a Regular Faculty Research Grant, Maine Top Scholar research supply funds, the Center for Undergraduate Research, the Frederick Radke Undergraduate Research Fellowship, and Charlie Slavin Research grants from the Honors College.

We thank Jessie Bruno, Emily Ledue, and Esther Biro for technical assistance and helpful scientific discussions.

## Declaration of Interest Statement

The authors declare that they have no conflicts of interest with the contents of this article.

## Notes

### Competing Interest Statement

The authors have declared no competing interest.

