## Supplement for "Antimicrobial cetylpyridinium chloride suppresses mast cell function by targeting tyrosine phosphorylation of Syk kinase"

**Table of Contents**

Determination of hexadecylbenzene (HDB) concentration………………………….…S1

CPC effects on constitutive tyrosine phosphorylation ………………………...………..S2

CPC effects on protein concentration in cell lysate ……………………………………. S3

Normalization to Revert 700 Total Protein Stain or β-actin…...………………………...S4

Linearity test to determine the right amount (µg) of protein for WB…………………… S5

**Determination of hexadecylbenzene (HDB) concentration**

**Methods**: HDB was prepared, as described in the main “Materials and Methods” section, in DMSO vehicle (0.046% final concentration) in an overall aqueous solution of Tyrodes buffer. The UV-Vis spectrum was collected before BSA was added.

**Results:** HDB absorbed maximally around 260 nm^1^ (Fig. S1). The concentration calculated via use of the CPC extinction coefficient, 10.2 µM, is ten-fold lower than the nominal value (106.6 µM, based on the grams of HDB added to the buffer and dilution factors).

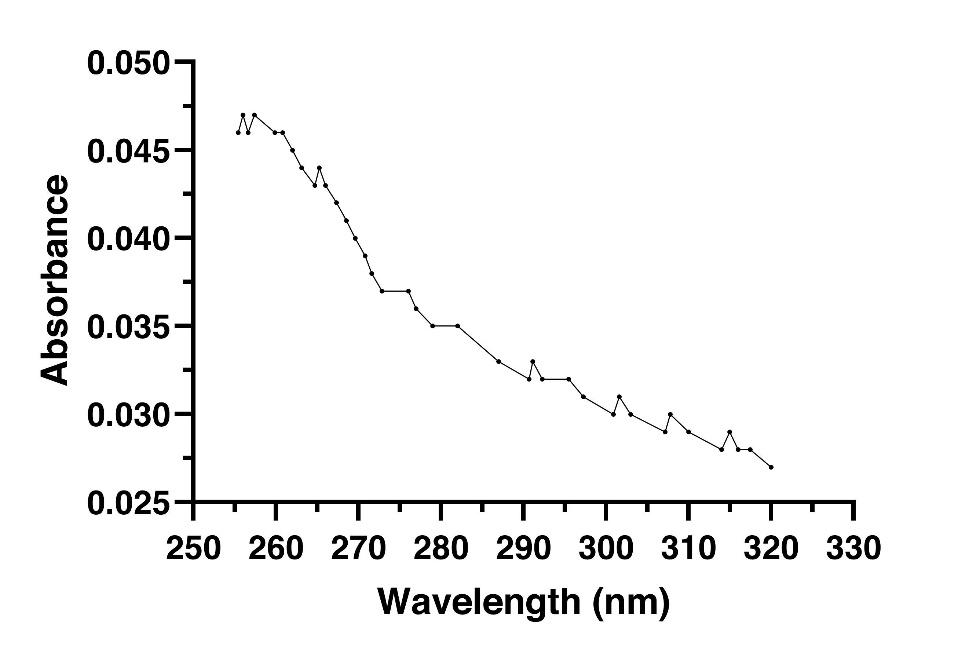

**Figure. S1. UV-Vis absorption spectrum of HDB**. HDB was first dissolved in 100% DMSO, sonicated, diluted in aqueous Tyrodes buffer, and absorption read using UV-Vis spectrophotometry.

**Conclusion**: HDB dissolves in an aqueous buffer, can be detected with UV-Vis spectrophotometry, and may be present at a significantly higher concentration than that calculated via CPC’s extinction coefficient.

**CPC effects on constitutive tyrosine phosphorylation**

An ICW experiment was performed to determine the effect of CPC on constitutive tyrosine phosphorylation in RBL mast cells.

**Method:** To assess the CPC effect on constitutive tyrosine phosphorylation (**Fig. S2**), the ICW procedure was followed as detailed in “Materials & Methods.” Cells were pre-treated with CPC (“Spont. + CPC”) or BT (“Spont.”) for 30 min. At the end of the pre-treatment, the ICW assay described in “Materials & Methods” was used to assess the CPC effect on constitutive phosphorylation. A unpaired, two-tailed Student’s t test was applied and revealed no significant difference.

**Result:** CPC does not affect constitutive tyrosine phosphorylation.

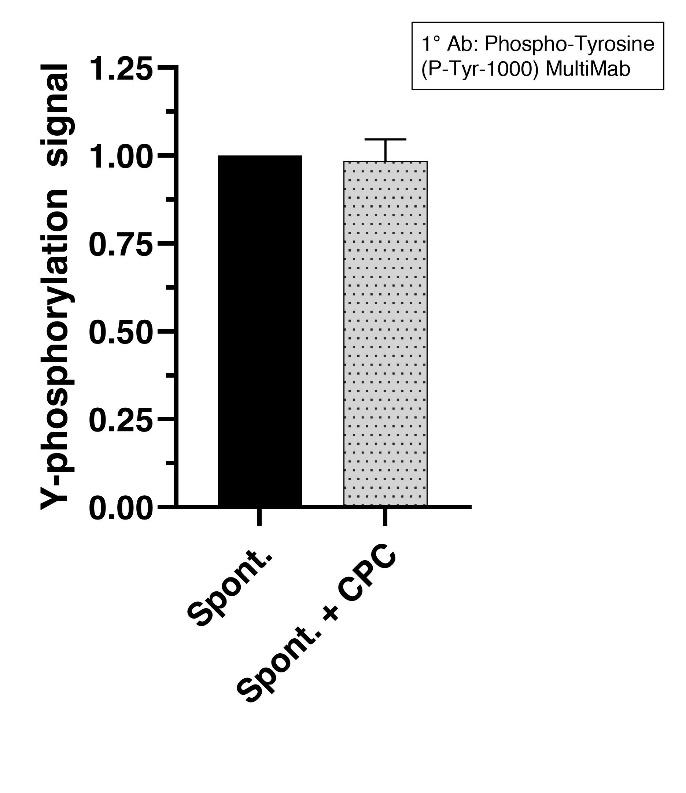
 Figure S2. **CPC effect on constitutive tyrosine phosphorylation** **assessed via ICW**. Values presented are means ± SD for triplicate measurements in a single representative experiment.

**Conclusion**: CPC does not affect constitutive tyrosine phosphorylation.

**CPC effects on protein concentration in cell lysate**

The effect of CPC on protein concentration in cell lysate following CPC treatment was assessed.

**Method:** The CPC effect on total protein concentration in equivalent wells of cells, following CPC pretreatment and Ag stimulation, was assessed as described in the “Materials and Methods” section. The mean protein concentration from the “Ag" and “Ag + CPC” groups for at least three independent days of the experiment was taken and compared via an unpaired two-tailed t-test.

**Results:** CPC does not affect the total protein concentration of the lysate in Ag-stimulated cells.

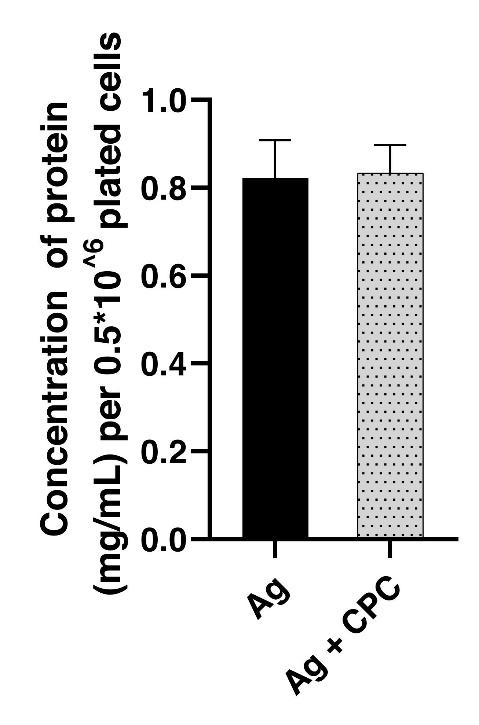

**Figure S3. CPC effect on protein concentration following treatment**. Cells were pretreated with CPC for 30 min and exposed to Ag ± CPC. Protein concentration was determined using RC DC protein assay as described in “Materials and Methods.”

**Conclusion:** CPC does not alter the total protein concentration during the western blot experimental workflow.

**Normalization to Revert**^TM^ **700 Total Protein Stain or β-actin**

Target proteins were normalized to total protein determined using Revert^TM^ 700 Total Protein Stain or β-actin.

**Method:** The Revert^TM^ 700 Total Protein Stain was performed according to the manufacturer’s instructions, as described in the “Materials and Methods” section**.** β-actin signal was determined as described in the “Materials and Methods” section. Normalization of bands of interest to Revert^TM^ 700 Total Protein Stain or β-actin was done as described in the “Materials and Methods” section.

**Results:** Protein transfer onto the membrane was efficient and consistent across all lanes.

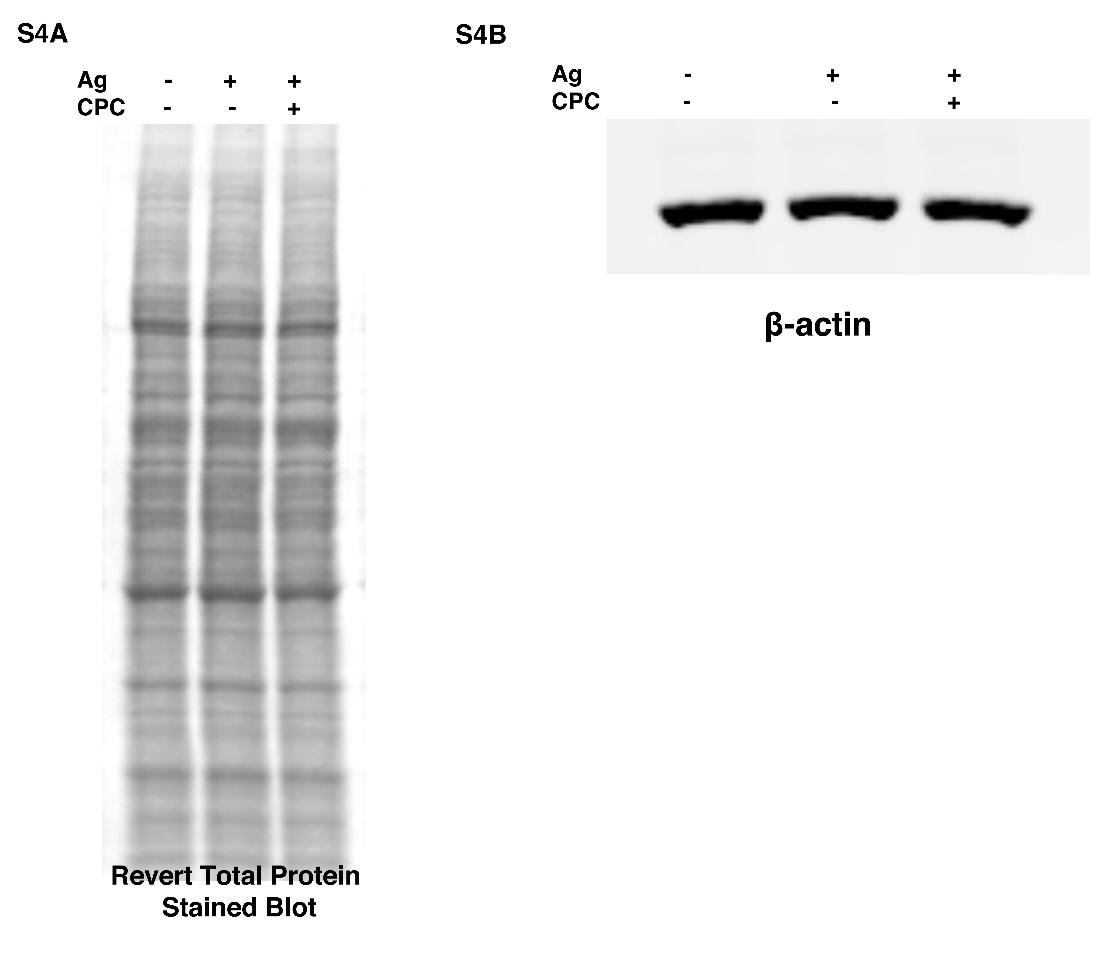

**Figure S4. Representative image of Revert^TM^ 700 Total Protein Stain (A) and β-actin (B).**

**Conclusion:** Whether Revert^TM^ 700 Total Protein Stain or β-actin was used, protein transfer onto the membrane was efficient and consistent across all lanes and, thus, suitable for normalization as described in Materials and Methods.

**Linearity test to determine the right amount (µg) of protein for WB**

A linearity test was done to determine the range within which WB signal from specific target proteins increased linearly with increasing amounts of loaded protein. The result was used to determine an appropriate amount of protein to load per lane, in order to be able to detect increases or decreases in the target due to CPC.

**Method:** RBL cells were Ag stimulated without CPC treatment, and western blotting was done as described in the “Materials and Methods” section, except that varying µg of proteins were loaded in individual lanes. After imaging, a curve with “mean fluorescence” on the Y-axis and µg of protein on the X for the proteins of interest was generated. The R^2^ value and the slope are determined. The R^2^ value is a measure of linearity. The slope represents the average intensity of LI-COR signal per µg of protein. After generating the curve, if the Y-intercept value (X = 0) was different from 0, then the Y-intercept was subtracted from the background-subtracted “mean fluorescence” values. Then a new curve was plotted with the Y-intercept-subtracted values. Subtracting the Y-intercept did not change the R^2^ values.

**Results**. The amount loaded, 12 µg, was within the linear range of all target proteins, β-actin, and total protein signal (determined using Revert 700 Total Protein Stain) with R-squared values ranging from 0.94 – 0.99 (indicated on individual curves in Fig S5)

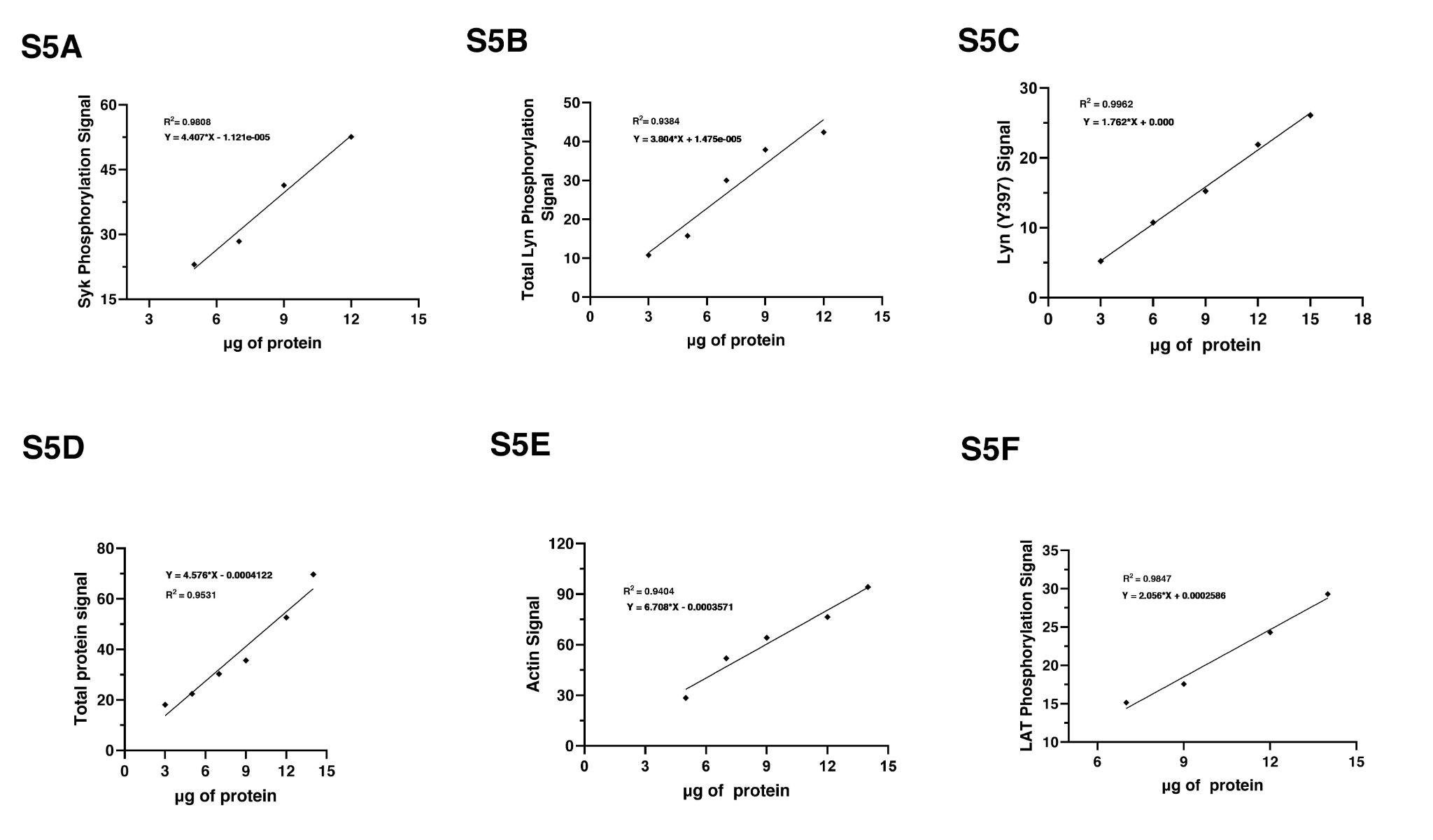

**Figure S5. Linearity test to determine the linear range of target proteins.**

**Conclusion:** The amount loaded, 12 µg, was within the linear range of all target proteins. R^2^ values ranged from 0.94 – 0.99.
